# Cell intrinsic mechanical regulation of plasma membrane accumulation at the cytokinetic furrow

**DOI:** 10.1101/2023.11.13.566882

**Authors:** Roberto Alonso-Matilla, Alice Lam, Teemu P. Miettinen

## Abstract

Cytokinesis is the process where the mother cell’s cytoplasm separates into daughter cells. This is driven by an actomyosin contractile ring that produces cortical contractility and drives cleavage furrow ingression, resulting in the formation of a thin intercellular bridge. While cytoskeletal reorganization during cytokinesis has been extensively studied, little is known about the spatiotemporal dynamics of the plasma membrane. Here, we image and model plasma membrane lipid and protein dynamics on the cell surface during leukemia cell cytokinesis. We reveal an extensive accumulation and folding of plasma membrane at the cleavage furrow and the intercellular bridge, accompanied by a depletion and unfolding of plasma membrane at the cell poles. These membrane dynamics are caused by two actomyosin-driven biophysical mechanisms: the radial constriction of the cleavage furrow causes local compression of the apparent cell surface area and accumulation of the plasma membrane at the furrow, while actomyosin cortical flows drag the plasma membrane towards the cell division plane as the furrow ingresses. The magnitude of these effects depends on the plasma membrane fluidity, cortex adhesion and cortical contractility. Overall, our work reveals cell intrinsic mechanical regulation of plasma membrane accumulation at the cleavage furrow that is likely to generate localized differences in membrane tension across the cytokinetic cell. This may locally alter endocytosis, exocytosis and mechanotransduction, while also serving as a self-protecting mechanism against cytokinesis failures that arise from high membrane tension at the intercellular bridge.

## MAIN TEXT

Cytokinesis, the process of partitioning a mother cell’s cytoplasm into daughter cells, is an essential step in the completion of the cell division cycle. The key biomechanical forces responsible for animal cell cytokinesis are generated by the actomyosin cortex, a network of myosin-II motors and crosslinked actin filaments (1). A localized increase in actomyosin contractility at the cell’s equatorial region, coupled with actomyosin relaxation at the cell poles generates a cortical contractility gradient that induces the movement of cortical material from the cell poles towards the equator (2, 3). At the cell’s equator, an actomyosin ring assembles, constricts, and forms a cleavage furrow, an indentation of the cell’s surface. The cleavage furrow, also known as the cytokinetic furrow, advances inwards until the daughter cells are only connected by a narrow intercellular bridge (4–6). These early stages of cytokinesis, which typically last only ∼10 min, are then followed by the formation of the midbody, reorganization of the cytoskeleton and eventually the final abscission of the intercellular bridge (5–7).

Cytokinetic fidelity is critical for maintaining cell fitness and tissue homeostasis, and for preventing cancer initiation and progression (8). Errors in cytokinesis can lead to imperfect partitioning of cellular components, and a failed cytokinesis will result in polyploidy or cell death. A successful completion of cytokinesis depends on the mechanical forces within the intercellular bridge. High tension in the intercellular bridge inhibits or delays cell abscission by hindering proper assembly and function of the abscission machinery. This is mediated, at least in part, by tension in the plasma membrane (7, 9–13), suggesting that cytokinetic fidelity would benefit from accumulation of plasma membrane at the cytokinetic furrow. However, except for exocytosis at the cytokinetic furrow that primarily occurs in late cytokinesis (14–20), little is known about the dynamics and the regulation of the plasma membrane during cytokinesis.

The plasma membrane is a highly folded fluidic structure that is predominantly composed of proteins (∼50%) and lipids (∼50%) and is attached to the actin cortex by membrane-cortex linker proteins (21). Several studies have shown that certain plasma membrane proteins and lipids accumulate at the cytokinetic furrow during cytokinesis. For example, caveolin-1, a key protein forming membrane reservoirs called caveolae, as well as several lipids, such as cholesterol and phosphoinositides, accumulate at the cytokinetic furrow and take part in the regulation of cytokinesis (9, 22–29). However, it remains unclear as to what extent the accumulation of specific membrane components to the cytokinetic furrow reflects the overall dynamics of the plasma membrane rather than a regulated localization of specific membrane components. Consequently, it is essential to gain a deeper understanding of how cytoskeletal forces regulate plasma membrane movement and tension at the cleavage furrow (30).

Here, we carry out live cell imaging and biophysical modeling of plasma membrane dynamics during leukemia cell cytokinesis. We discover that the plasma membrane accumulates at the cytokinetic furrow, and we reveal two cell-intrinsic biophysical mechanisms responsible for this behavior. Our results show that cytoskeletal mechanisms driving cell division also act to decrease membrane tension at the cleavage furrow and intercellular bridge, thereby assisting in the completion of cytokinesis.

## RESULTS

### Global accumulation of plasma membrane proteins at the cleavage furrow and intercellular bridge

We sought to study how cell-intrinsic factors regulate plasma membrane movement on the cell surface. We labeled all plasma membrane proteins accessible on the external side of the cell using an amine-reactive, cell-impermeable, fluorescent label (Supplementary Note 1). As a model system, we used a suspension grown mouse lymphocytic leukemia cell line, L1210, that expressed histone H2B fused with green fluorescent protein (H2B-GFP) to visualize DNA. These cells are spherical and display little to no attachment to the growth substrate or neighboring cells. Cells in interphase and early mitosis (prophase to metaphase) displayed an even distribution of plasma membrane proteins on all sides of the cell (Fig. 1A). However, during early cytokinesis, the plasma membrane protein content polarized, with proteins accumulating at the cytokinetic furrow (Fig. 1A). This membrane protein accumulation was initially evident in anaphase cells, coinciding with the onset of cleavage furrow constriction, and the protein accumulation increased with the progression of cytokinesis (Fig. 1B, Fig. S1). Later in cytokinesis, membrane protein accumulation was predominantly observed in the intercellular bridge (Fig. 1A, B). Similar membrane protein distributions were seen across all L1210 cells imaged (Fig. 1C), and when using different labeling chemistry (Fig. S2H).

**Figure 1.**
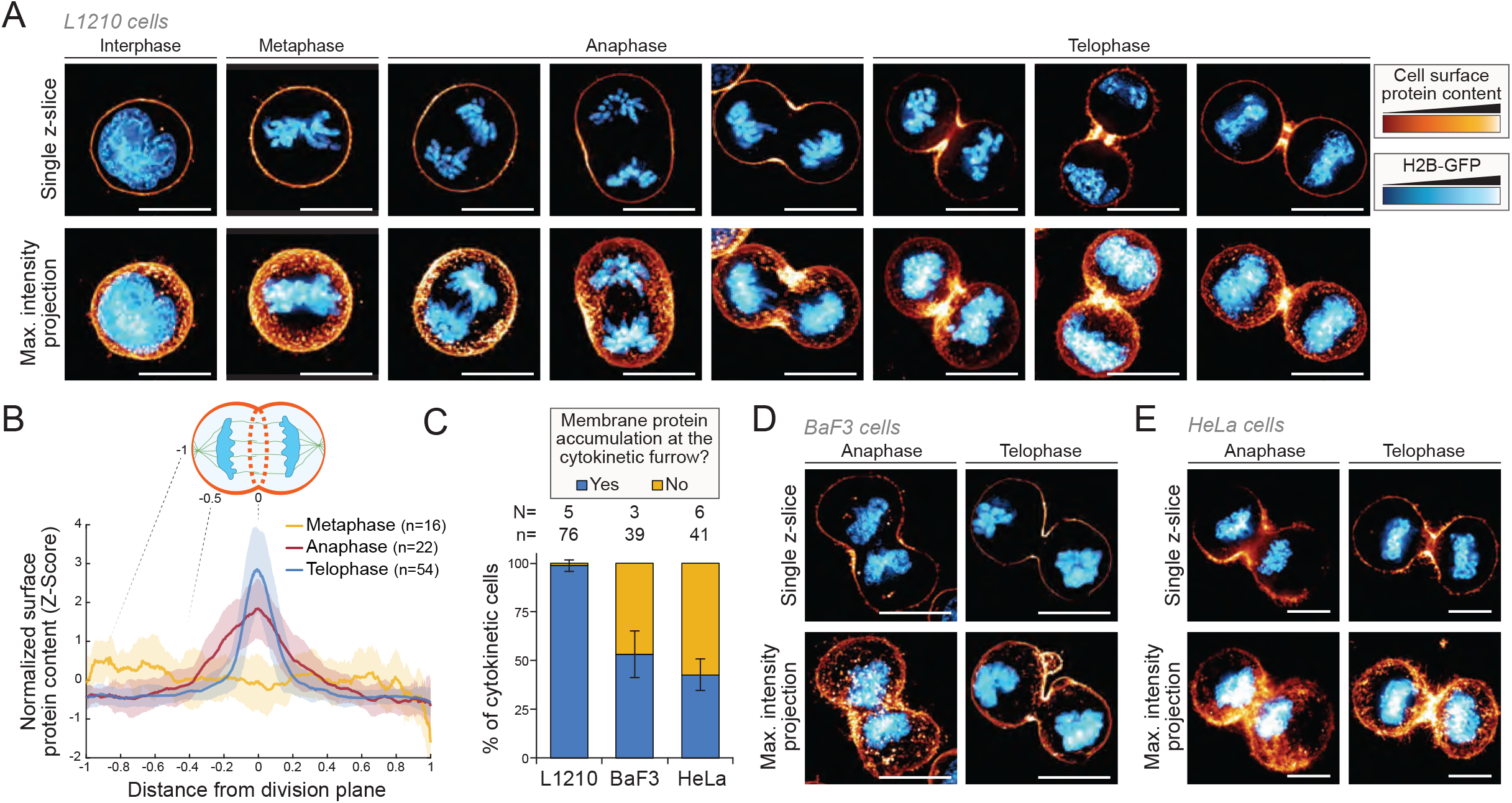
Cell surface proteins accumulate at the cytokinetic furrow and intercellular bridge. **(A)** Representative images of mitotic and cytokinetic L1210 cells expressing H2B-GFP (blue, DNA) and labeled for cell surface protein content (orange/yellow). Cells were imaged immediately after labeling (i.e., late cytokinetic cells were labeled in late cytokinesis). **(B)** Quantifications of cell surface protein content as a function of distance from the division plane, as indicated in the schematic on top. Lines and shaded areas depict mean±SD, n depicts the number of individual cells in each cell cycle stage. **(C)** Percentage of cytokinetic cells that exhibit the accumulation of plasma membrane proteins at the cleavage furrow in the indicated cell lines. Data depicts mean±SD of independent experiments. N depicts the number of independent experiments and n depicts the total number of cytokinetic cells. **(D, E)** Representative images of cytokinetic BaF3 (D) and Hela (E) cells expressing H2B-GFP (blue) and labeled for cell surface protein content (orange/yellow). All scale bars denote 10 μm.

To ensure that these plasma membrane protein dynamics were not limited to a single cell type, we imaged a suspension grown mouse pro-B cell myeloma cell line (BaF3), and an adherent human adenocarcinoma cell line (HeLa). Both cell lines displayed an accumulation of plasma membrane proteins at the cleavage furrow (Fig. 1D, E), although this effect was not uniform across cells (Fig. 1C, Supplementary Note 2). As all three cell lines displayed the capacity to accumulate membrane at the cytokinetic furrow, from here on we investigated plasma membrane dynamics using L1210 cells as our primary model system.

### Plasma membrane folds at the cleavage furrow and unfolds at the cell poles

We considered two hypothetical models of plasma membrane composition and structure that could explain the high membrane protein content at the cleavage furrow (Fig. 2A). In model 1, the surface density of proteins, i.e., the number of proteins per unit of membrane area, at the cleavage furrow increases. In this model, plasma membrane lipids do not accumulate at the furrow. In model 2, the surface density of proteins remains constant, and plasma membrane area increases at the furrow due to membrane folding (Fig. 2A). This would result in plasma membrane lipids, carbohydrates, and other membrane components displaying a higher density at the furrow. Importantly, in model 2, plasma membrane accumulation at the cleavage furrow predicts a local decrease in membrane tension at the furrow, whereas model 1 would not result in significant membrane tension changes at the furrow.

**Figure 2.**
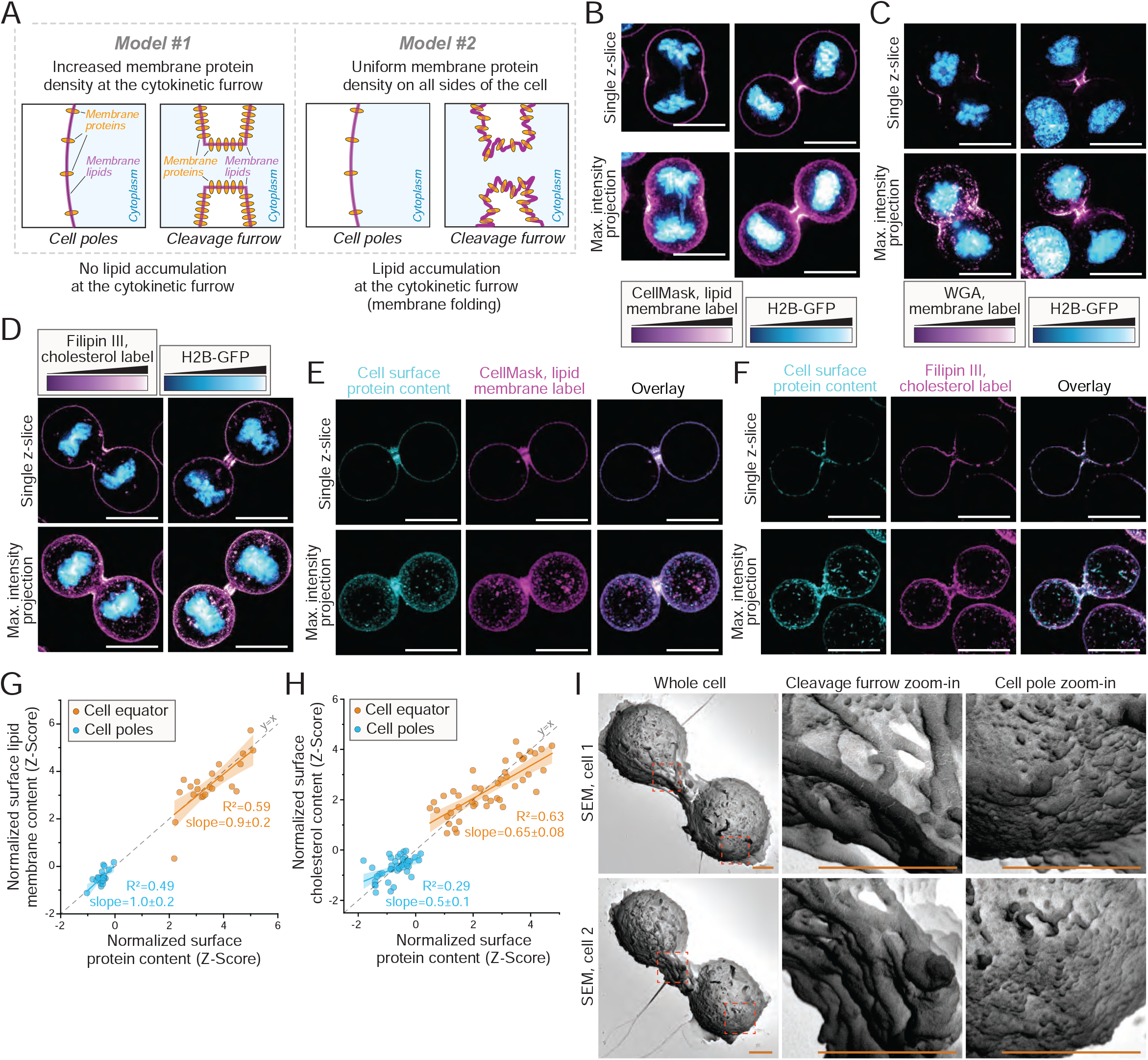
Excess plasma membrane accumulates at the cytokinetic furrow and depletes at the cell poles. **(A)** Two competing models that can explain the accumulation of plasma membrane proteins at the cytokinetic furrow. **(B-D)** Representative images of cytokinetic L1210 cells expressing H2B-GFP (blue, DNA) and labeled for plasma membrane (purple/white) using CellMask lipid label (panel B, N=3 independent experiments, n=46 cytokinetic cells), WGA (panel C, N=2 independent experiments, n=31 cytokinetic cells), or Filipin III (panel D, N=3 independent experiments, n=68 cytokinetic cells). **(E-F)** Representative images of cytokinetic L1210 cells labeled using cell surface protein label (teal) together with CellMask lipid label (purple) (E) or Filipin III cholesterol label (F) (N=4 or 3 independent experiments and n=22 or 44 cytokinetic cells, for panels F and E respectively). **(G-H)** Correlation between cell surface lipid (G) or cholesterol (H) content and protein content at cell poles (light blue) and cell equator (orange). Data is the same as in panels E-F. Dots depict single cells, lines and shaded areas depict linear fits and their 95% confidence intervals. Slope error values depict 95% confidence intervals. **(I)** Representative SEM images of L1210 cells in cytokinesis (N=3 independent experiments, n=45 cytokinetic cells). Fluorescence imaging scale bars denote 10 µm and SEM scale bars denote 2 μm.

To distinguish these two models, we first labeled live L1210 cells with a CellMask plasma membrane lipid stain. This revealed that plasma membrane lipids accumulate at the cytokinetic furrow and the intercellular bridge (Fig. 2B, Fig. S3A), consistent with previous observations in *Drosophila* neuroblasts (31). Approximately 60% of BaF3 cells also displayed accumulation of membrane lipids at the cytokinetic furrow (Fig. S4A,B,E). Then, we labeled live L1210 cell plasma membrane using wheat germ agglutinin (WGA), which binds carbohydrates on glycoproteins and glycolipids. WGA labeling accumulated to the cytokinetic furrow and intercellular bridge (Fig. 2C, Fig. S3B). We also labeled plasma membrane cholesterol in fixed cells using Filipin III, which revealed cholesterol accumulation at the cytokinetic furrow (Fig. 2D, Fig. S3C), consistent with previous reports in various model systems (27–29). With all three approaches, we observed that plasma membrane was distributed evenly on all sides of the cell prior to cytokinesis.

We then co-labeled membrane lipids and cell surface proteins in live L1210 cells (Fig. 2E, Fig. S3D) and examined the correlation between cell surface lipid and protein content in cytokinetic cells. This revealed that lipid and protein components within the plasma membrane accumulate and deplete similarly to each other at the cell equator and poles, respectively (Fig. 2G). When we co-labeled L1210 cells for cholesterol and cell surface proteins (Fig. 2F), however, we observed that although both components accumulate at the cytokinetic furrow and deplete at cell poles, these effects are more pronounced for cell surface proteins than for cholesterol (Fig. 2H). Overall, these results support our model 2 (Fig. 2A), wherein the plasma membrane protein accumulation at the cytokinetic furrow is mostly attributed to a local increase in cell surface area (i.e., membrane folding).

To verify that plasma membrane reservoirs accumulate at the cytokinetic furrow, we imaged L1210 cells using scanning electron microscopy (SEM). We observed that the plasma membrane at the cytokinetic furrow displays significantly greater structural complexity and membrane folds compared to the membrane observed at the cell poles (Fig. 2I, Fig. S5A). This difference was particularly pronounced during early cytokinesis, where the cytokinetic furrow and intercellular bridge displayed large membrane folds. This membrane accumulation was specific to cytokinesis, as cells arrested in prometaphase by the kinesin inhibitor S-trityl-l-cysteine (STLC) displayed uniform membrane folding on all sides of the cell (Fig. S5B). As the cytokinetic furrow fully closed, the membrane accumulation became undetectable in SEM images (Fig. S5A). We also carried out SEM imaging of cytokinetic BaF3 cells. These cells displayed additional membrane folding at the cleavage furrow (Fig. S5C), although the membrane accumulation was not observed in every cell. Overall, we conclude that the plasma membrane, rather than particular membrane components, accumulates at the cytokinetic furrow. Consequently, our results suggest the existence of membrane tension gradients between the cell equator (low tension) and cell poles (high tension) during cytokinesis.

### Null models for the biophysical basis of plasma membrane dynamics during cytokinesis

We then investigated the biophysical mechanism(s) responsible for membrane accumulation at the cytokinetic furrow. We considered three potential mechanisms that can cause local membrane accumulation (or depletion): First, actomyosin driven changes in cell shape during cytokinesis can locally alter the apparent cell surface area which could influence plasma membrane accumulation (Fig. 3A). Second, cortical flows, driven by cortical contractility gradients, are directed towards the cell division plane (2, 3), and membrane-cortex drag forces (33–37) could pull the plasma membrane towards the cleavage furrow (26, 31, 35) (Fig. 3A). Third, endocytosis and exocytosis can locally decrease and increase plasma membrane content, respectively. If exocytosis is localized to the cleavage furrow and endocytosis to the cell poles during early cytokinesis, then this membrane trafficking could increase plasma membrane area at the furrow (Fig. 3A). These three mechanisms are not mutually exclusive, and each of them could cause plasma membrane to accumulate at the cytokinetic furrow while depleting plasma membrane at the cell poles, as observed in our experiments.

**Figure 3.**
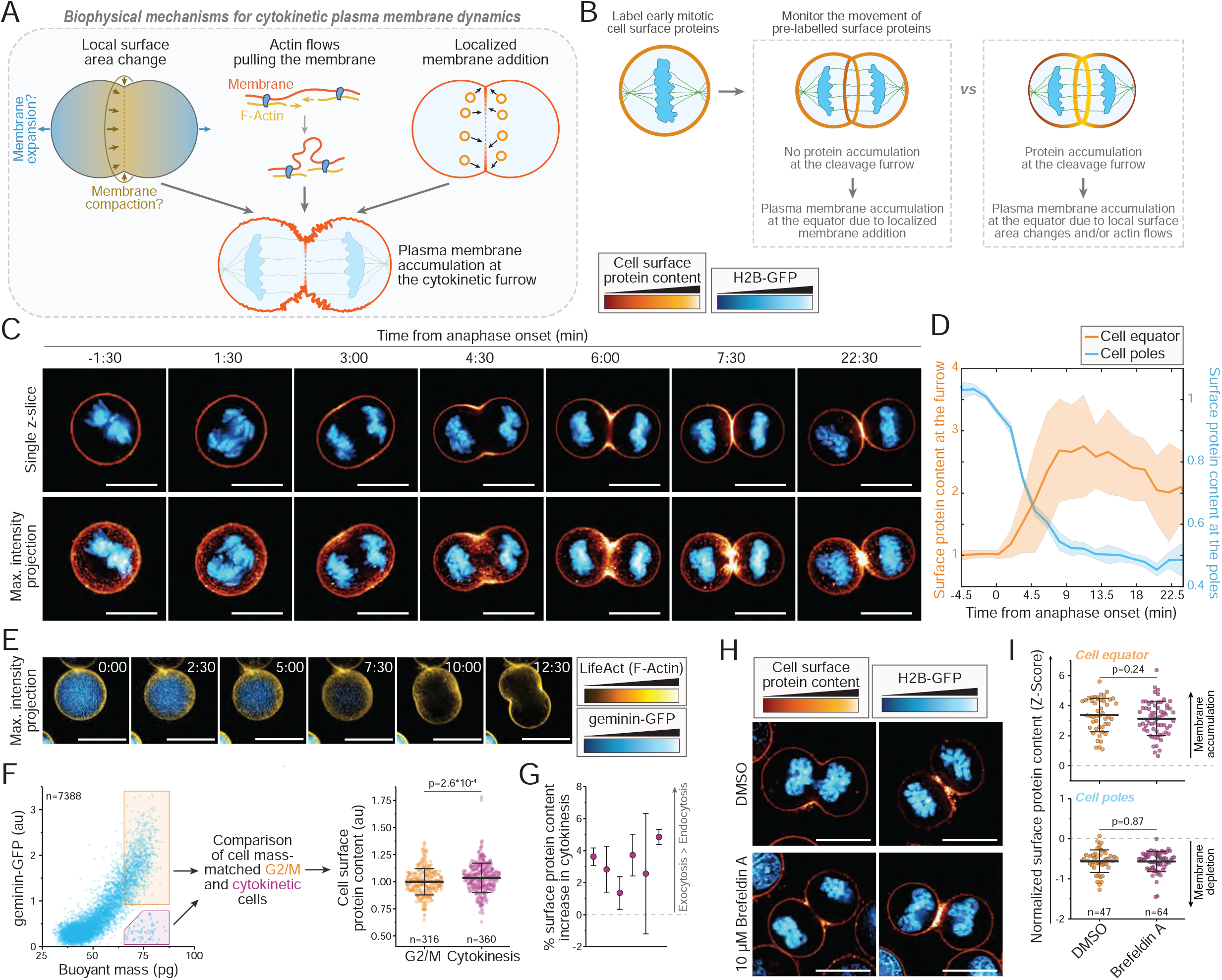
Plasma membrane accumulation at the cleavage furrow is caused by movement of plasma membrane reservoirs on cell surface. **(A)** Three mutually non-exclusive biophysical processes that can explain the plasma membrane accumulation at the cytokinetic furrow. **(B)** Experimental setup: Pre-labeled plasma membrane proteins are imaged over time to examine whether the pre-labeled proteins accumulate at the cytokinetic furrow. Since internal membranes are not labeled, exocytosis-driven membrane accumulation will not result in a detectable signal accumulation. **(C)** Representative timelapse imaging of L1210 cells expressing H2B-GFP and labeled for cell surface protein content (orange/yellow). All cells were labeled before the onset of cytokinesis. **(D)** Quantifications of L1210 cell surface protein content at the cytokinetic furrow (orange) and cell poles (blue) as a function of time from anaphase onset. Lines and shaded areas depict mean±SD (N=3 independent experiments, n=14 cells). Data is normalized to the values at t≤0. **(E)** Example timelapse images of an L1210 cell expressing geminin-GFP (blue) and LifeAct (yellow). Geminin-GFP is rapidly degraded at the onset of anaphase. Time (white text) is in minutes. **(F)** Experimental setup: *Left*, geminin-GFP expressing cells were labeled for surface protein content and analyzed using the SMR, so that cytokinetic cells can be compared to equally sized G2/M cells. *Right*, pooled surface protein labeling data for L1210 cells. Lines and whiskers depict mean±SD, dots depict individual cells, n depicts the number of cells (N=6 independent experiments). **(G)** % increase in cell surface protein content of cytokinetic cells in comparison to equally sized G2/M cells. Each dot is an independent experiment, whiskers depict SEM. **(H)** Representative single z-layer images of cytokinetic L1210 cells expressing H2B-GFP (blue) and labeled for surface protein content (orange/yellow) after a 1 h treatment with DMSO or 10 µM Brefeldin A. **(I)** Quantifications of the surface protein content at the equatorial and polar regions in the samples shown in panel (H). Lines and whiskers depict mean±SD, dots depict individual cells, n depicts the number of cells (N=3 independent experiments). All p-values were obtained using Welch’s t-test. All scale bars denote 10 µm.

### Plasma membrane accumulation at the cleavage furrow is caused by movement of plasma membrane reservoirs on the cell surface

To identify the biophysical mechanism(s) responsible for plasma membrane accumulation at the cytokinetic furrow, we first examined if the membrane accumulation can be explained by the movement of plasma membrane on the cell surface or if exocytosis is required. We labeled plasma membrane surface proteins in early mitosis and tracked these pre-labeled surface proteins (Fig. 3B). If plasma membrane accumulation at the cleavage furrow is driven primarily by localized insertion of internal vesicles, i.e., exocytosis, we would not observe membrane accumulation, as the newly added membrane was not pre-labeled. However, if the plasma membrane accumulation is primarily driven by membrane movement on the cell surface, then we should observe the pre-labeled membrane accumulate at the furrow (Fig. 3B). Live cell imaging of both L1210 and HeLa cells indicated that the pre-labeled membrane proteins accumulate at the cleavage furrow and deplete at the cell poles as cytokinesis progresses (Fig. 3C, Fig. S6, Movie S1). In L1210 cells, the surface protein accumulation at the cleavage furrow reached its maximum within 10 minutes of cytokinesis, as the furrow ingression completed (Fig. 3D). In later cytokinesis, the surface protein accumulation slowly dissipated. Intracellular fluorescence signals remained consistently low throughout the timelapse imaging, especially in L1210 cells, indicating minimal endocytosis of the labeled plasma membrane proteins. Thus, plasma membrane reservoirs move on the cell surface towards the cleavage furrow during cytokinesis, and the plasma membrane accumulation partially persists to late cytokinesis.

Exocytosis is known to be active in cytokinesis, but the magnitude of cytokinetic exocytosis, especially in comparison to endocytosis, remains poorly defined. To study this, we examined changes in plasma membrane protein content that are specific to cytokinetic cell state and independent of cell growth. We labeled cell surface proteins in cells expressing geminin-GFP cell cycle indicator which is rapidly degraded at the onset of anaphase (Fig. 3E) (38). In L1210 cells, geminin-GFP degradation takes only 9 minutes (39). We then measured these cells using the suspended microchannel resonator (SMR), which is a single-cell mass sensor coupled with fluorescent detection setup (39–41). This allowed us to compare the surface protein content of cells in G2/M phase with the surface protein content of identically sized cells in cytokinesis (Fig. 3F). We found that L1210 cells have a higher plasma membrane protein content in cytokinesis than in G2/M, but the increase was only 3% on average (Fig. 3G). Thus, we can rule out overall increases in exocytosis (or overall decreases in endocytosis) as the cause of membrane accumulation at the cytokinetic furrow, although localized endocytosis and exocytosis may still contribute to the membrane accumulation at the furrow.

To further examine the role of localized exocytosis in membrane accumulation at the cytokinetic furrow, we treated L1210 cells with Brefeldin A, an inhibitor of endocytosis and exocytosis (42) which significantly decreases exocytosis in L1210 cells without preventing cell division (40). Brefeldin A did not prevent the accumulation of plasma membrane at the equatorial region nor the depletion of plasma membrane at cell poles (Fig. 3H,I). Overall, these results suggest that the accumulation of plasma membrane at the cleavage furrow can be predominantly attributed to the movement of membrane reservoirs on the cell surface.

### A force balance model of cytokinesis

Next, we aimed to investigate how local changes in cell surface area and actin flows impact the accumulation of plasma membrane components at the cytokinetic furrow. To uncouple the individual impacts of these two factors, we opted for a modeling approach. By building on previous theoretical work (43, 44), we developed a three-dimensional biophysical model of actomyosin-driven cytokinesis, that includes the hydrodynamics of lipid flows during furrow ingression as a fundamental component (Fig. 4A, Supplementary Note 3). The model considers the cellular cortex as a thin actomyosin film that produces force and creates stresses in the cortical network (1, 45–47), and enforces force balance and mass conservation of cortex components, specifically actin and myosin (Fig. 4B).

**Figure 4.**
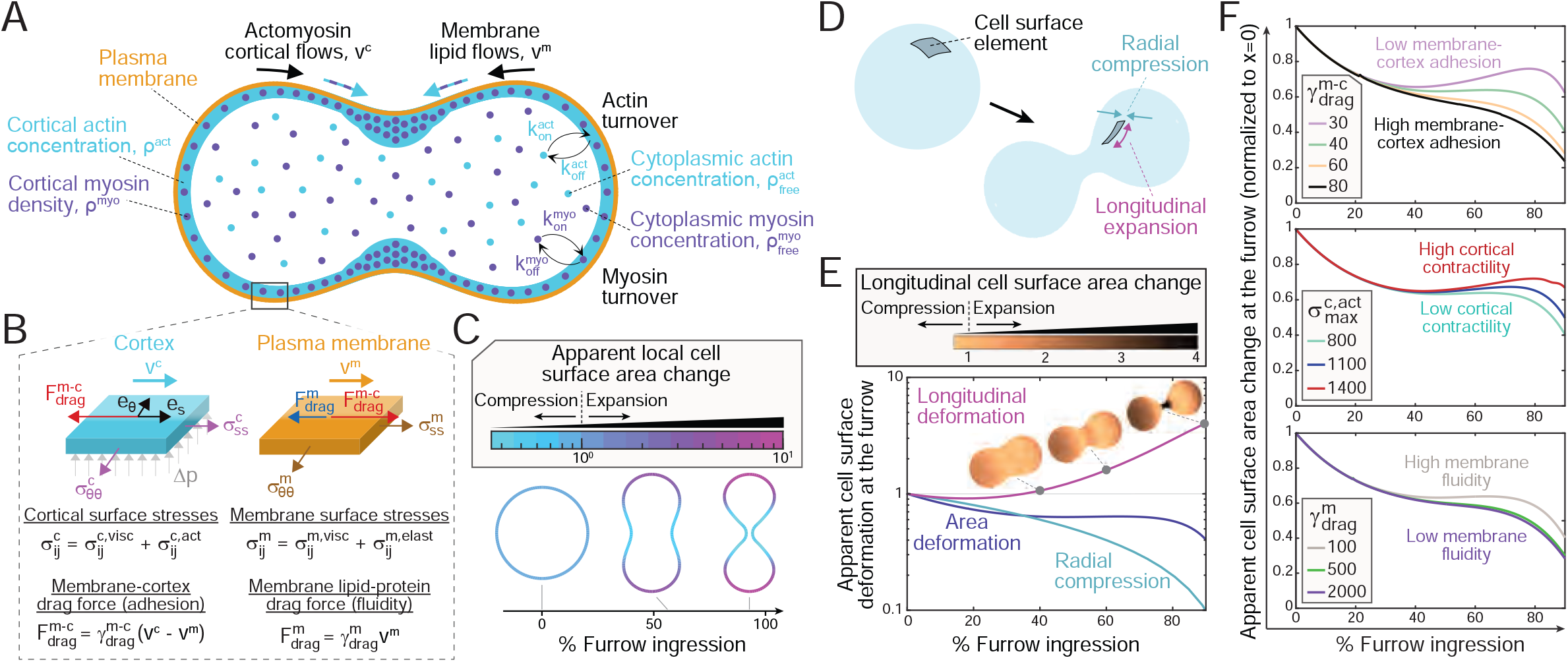
A force balance model of cytokinesis reveals local cell surface area changes due to radial constriction and longitudinal stretching. **(A)** Schematic of the biophysical model of cytokinesis. Force balance, transport of cortical components and hydrodynamics of lipid flows during furrow ingression are fundamental components of the model. **(B)** Surface stresses and body forces on infinitesimal cortex and plasma membrane surface elements. Cortical stress forces are balanced by pressure forces and membrane-cortex adhesion forces. The cortical stress is equal to the sum of a viscous contribution 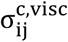 and an active contribution 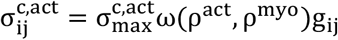, where 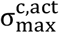 is cortical contractility, ωis a cortex density-dependent saturating function, and i and j indicate the components of the stress tensor. Plasma membrane stress forces are balanced by drag forces. The membrane stress is the sum of viscous and elastic stresses. **(C)** Local changes in apparent cell surface area at three different timepoints during cytokinesis. **(D)** Schematic of the local changes in the apparent cell surface area due to radial and longitudinal deformations. **(E)** Apparent cell surface deformation at the cytokinetic furrow as furrow ingression progresses. Data is shown separately for apparent total area (navy), radial compression (teal), and longitudinal deformation (light purple). The cells on top depict the degree of longitudinal deformation at different locations on cell surface. **(F)** Apparent cell surface area changes at the furrow during furrow ingression for different levels of membrane-cortex adhesion (displayed in units of pN ·s ·μm^−3^), cortical contractility (displayed in units of pN·μm^−1^), and membrane lipid-protein drag coefficient (displayed in units of pN·s·μm^−3^). Unless stated otherwise, model parameter values used are: 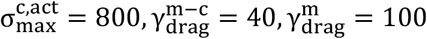.

Gradients in cortical contractility 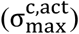 drive cortical flows and cell shape changes. We assume that the cortical network behaves as an active viscous material that undergoes fast rearrangements and dissipating stresses. The cytoskeletal network is subjected to membrane-cortex adhesion, as defined by an effective membrane-cortex drag coefficient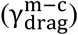, resulting in drag forces that slow down the movement of cortex constituents. The plasma membrane behaves as a two-dimensional viscous fluid with elastic properties. Plasma membrane lipid movement is caused by cortical flows via mobile membrane-cortex protein linkers and resisted by other proteins embedded within the plasma membrane (43). We characterize the fluidity of the plasma membrane by an effective membrane lipid-protein drag coefficient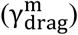, so that the membrane fluidity is inversely associated with the drag coefficient.

Cell shape is set by the cortical configuration. Plasma membrane lipid content is quantified as a projection of lipids onto the apparent cell surface, so that a local membrane accumulation (i.e., folding) is reflected as an increased lipid density. We used experimental data from the literature to parameterize our biophysical model and numerically solved the model equations (Supplementary Note 3). Our model captured the accumulation of cortical components at the division site (Fig. S7) and reproduced the approximate cell shapes and F-actin dynamics observed in L1210 cells expressing the LifeAct F-actin sensor (Fig. S8). Thus, our model mimics the cell-intrinsic regulation of actomyosin observed during cytokinesis, thereby enabling us to investigate how the mechanical properties of the actin cortex and plasma membrane impact membrane accumulation at the furrow.

### Local cell surface area compression can accumulate plasma membrane at the cytokinetic furrow

Using our biophysical model, we first examined local changes in cell shape and surface area during furrow ingression. By the end of furrow ingression, the total apparent cell surface area increased ∼30% (Fig. S9). The apparent cell surface area locally compressed up to ∼2.5-fold at the cytokinetic furrow and expanded up to ∼10-fold at the cell poles (Fig. 4C,D, Movies S2, S3). Consequently, furrow ingression alone can induce significant plasma membrane folding at the furrow, as driven by localized surface area compression, while simultaneously causing membrane unfolding at the cell poles due to local surface area expansion. We then analyzed radial and longitudinal cell surface area deformations at the cell’s equator in more detail. At the onset of furrow ingression, the furrow experienced weak longitudinal compression. As furrow ingression progressed further, the furrow experienced longitudinal stretching of up to ∼3-fold (Fig. 4E, Movies S2, S3). However, circumferential compression dominated over longitudinal stretching, causing an overall cell surface area compression at the furrow.

Our model enabled us to examine the mechanical factors that influence the apparent cell surface area changes at the cleavage furrow. Under high membrane-cortex adhesion, low cortical contractility, or low membrane fluidity conditions, longitudinal stretching was notably weaker than radial compression throughout furrow ingression, decreasing the apparent cell surface area at the furrow (Fig. 4F). In contrast, cells with high cortical contractility or weak membrane-cortex adhesion exhibited stronger cortical flows directed towards the cell’s equator, resulting in increased accumulation of cortical material and elevated contractility at the furrow (Fig. S7). This exacerbated longitudinal stretching of the cell surface at the furrow, reducing the overall local compression of the apparent cell surface area at the furrow (Fig. 4F, and Fig. S10A). Overall, our results show that the apparent cell surface area experiences significant compression irrespective of the mechanical factors responsible for shaping the cell. Thus, local surface area compression promotes plasma membrane accumulation at the furrow.

### Actomyosin-driven membrane flows and local surface area constriction are both responsible for the plasma membrane accumulation at the cleavage furrow

Our force balance model of cytokinesis captures the plasma membrane accumulation at the cleavage furrow (Movies S4, S5). Mechanistically, we found that high membrane fluidity and high membrane-cortex adhesion enhanced the accumulation of plasma membrane at the cleavage furrow (Fig. 5A). Interestingly, cortical contractility exhibited minimal impact on plasma membrane accumulation in cells with fluid-like membranes (Fig. 5A). In cells with low membrane drag coefficient, elevated cortical contractility resulted in a reduction in the apparent compression of cell surface at the furrow, but its effect on plasma membrane accumulation was largely cancelled out by strengthened pole-to-equator cortical flows, which also contribute to membrane accumulation at the furrow. In contrast, high cortical contractility significantly decreased plasma membrane accumulation at the furrow in cells with gel-like membranes (Fig. S10B-E). Under these conditions, the accumulation of plasma membrane at the cleavage furrow is near-exclusively dictated by changes in cell surface area. Notably, our model suggests that cells with low membrane fluidity, low membrane-cortex adhesion and high cortical contractility do not exhibit plasma membrane accumulation at the cytokinetic furrow (see Supplementary Note 2 and Fig. S10).

**Figure 5.**
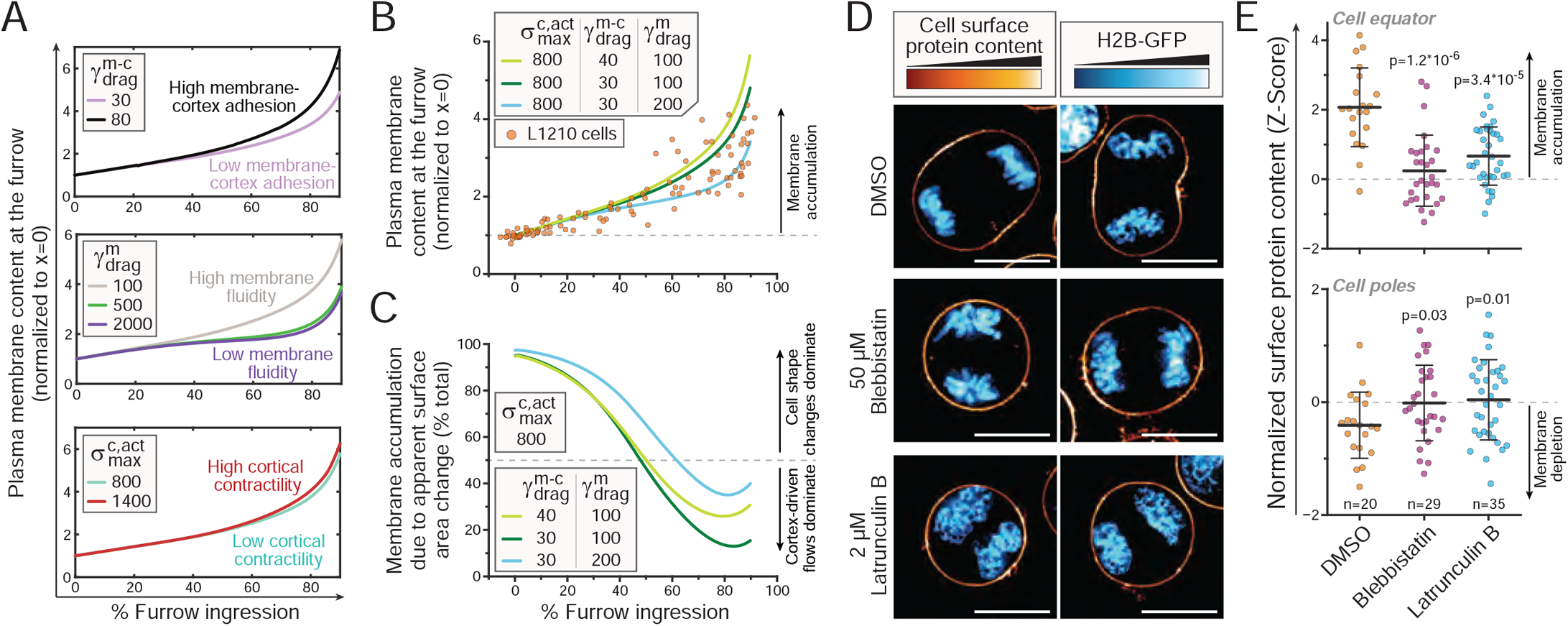
Actomyosin-driven membrane flows and local surface area constriction are both responsible for the plasma membrane accumulation at the cleavage furrow. **(A)** Predicted plasma membrane accumulation at the furrow as furrow ingression progresses for different values of the membrane-cortex drag coefficient 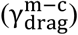, membrane lipid-protein drag coefficient 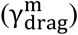, and cortical contractility 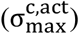. **(B)** Comparison of plasma membrane accumulation at the cytokinetic furrow in the force balance model and in live cell timelapse imaging as furrow ingression progresses. Lines depict modeling results using indicated model parameter values. Each orange dot depicts a cell at a given moment (n=14 cells with total of 103 images). **(C)** Relative contribution of apparent cell surface area changes, in comparison to cortex driven membrane flows, to the accumulation of plasma membrane at the cleavage furrow as furrow ingression progresses. Unless stated otherwise, model parameter values used in panels (A-C) are: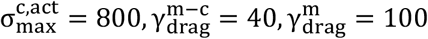. **(D)** Representative images of L1210 cells expressing H2B-GFP (blue) and labeled for surface protein content (orange/yellow) after a 2 h treatment with DMSO, 50 µM (-)-Blebbistatin, or 2 µM Latrunculin B. Scale bars denote 10 µm. **(E)** Quantifications of the surface protein content at the equatorial and polar regions in the samples shown in panel (D). Lines and whiskers depict mean±SD, dots depict individual cells, n depicts the number of cells (N=2-4 independent experiments), and p-values depict comparisons to DMSO (Welch’s t-test).

Importantly, the modelled plasma membrane accumulation at the cleavage furrow reproduced the membrane accumulation observed in live cells (Fig. 5B), for a cortical contractility of 800pN·μm^−1^, membrane-cortex drag coefficient of 30pN·s·μm^−3^, and membrane lipid-protein drag coefficients of 100−200pN·s·μm^−3^. These modeling parameters are in line with live cell measurements (45, 48–50). With these modeling parameters, the accumulation of plasma membrane at the cytokinetic furrow is not sensitive to changes in cortical contractility (Fig. 5A). Consistently, chemical perturbations of cortical contractility in L1210 cells did not alter plasma membrane accumulation at the cytokinetic furrow (see end of Supplementary Note 3, Fig. S11). As exocytosis is not included in our modeling approach, these results indicate that the plasma membrane accumulation at the cleavage furrow in our timelapse imaging can be explained solely by local cell surface area changes and cortex-driven membrane flows.

We then examined the relative impact that local cell surface area changes and cortex-driven membrane flows have on the accumulation of plasma membrane at the cytokinetic furrow. Using modeling parameter values that capture the membrane accumulation in live cells, we found that the local cell surface area changes and the cortex-driven membrane flows are approximately equally responsible for the membrane accumulation at the furrow (Fig. 5C). At the beginning of cleavage furrow ingression, plasma membrane accumulation at the furrow was dominated by local surface area changes, whereas cortical flows contributed more as the furrow ingression exceeded ∼50% ingression (Fig. 5C). However, the degree to which local cell surface area changes were responsible for the accumulation of plasma membrane at the cytokinetic furrow was dependent on the fluidity and cortex adhesion of the plasma membrane. Local cell surface area changes dominated plasma membrane accumulation at the furrow in cells with a gel-like plasma membrane and high membrane cortex adhesion, while cortical flows emerged as the primary driving force for plasma membrane accumulation in cells with more fluid-like plasma membranes and low membrane cortex adhesion (Fig. S12A, C). Cortical contractility had only a minor influence on the relative contribution of local cell surface area changes to plasma membrane accumulation at the furrow (Fig. S12B).

Overall, we conclude from our modeling that plasma membrane fluidity, membrane-cortex contacts, and cortical contractility define the plasma membrane accumulation at the cleavage furrow and the predominant mechanism responsible for the membrane accumulation. Importantly, the impact of cortical contractility and membrane-cortex adhesion on the plasma membrane accumulation was dependent on the fluidity of the plasma membrane. Consequently, cells may be able to regulate their membrane accumulation at the cytokinetic furrow by adjusting the mechanical properties of the plasma membrane and actin cortex. Additional regulation of the plasma membrane dynamics by anisotropic cortical contractility is detailed in Supplementary Note 4.

Finally, we validated the actomyosin-dependence of the plasma membrane dynamics. We treated L1210 cells with (-)-Blebbistatin, a non-muscle myosin II inhibitor, or Latrunculin B, an actin polymerization inhibitor, and examined plasma membrane and F-actin distributions. Treatment with both Blebbistatin and Latrunculin B prevented cytokinetic furrow ingression, the accumulation of plasma membrane and F-actin at the equatorial region, and the depletion of plasma membrane and F-actin at cell poles (Fig. 5D,E, Fig. S13). Thus, the accumulation of plasma membrane at the cytokinetic furrow is contingent upon actomyosin activity. In addition, Latrunculin B treatment caused F-actin to accumulate at random locations (Fig. S13), whereas plasma membrane protein content did not display as strong random accumulation (Fig. 5D). As Latrunculin B prevents actin polymerization, the random F-actin localization is primarily driven by cortex flows. Therefore, a local accumulation of F-actin by cortex flows is not always sufficient to drive extensive membrane accumulation at the same location.

## DISCUSSION

During cytokinesis, cells undergo major mechanical deformations that result in significant structural alterations to the plasma membrane. Here, we have revealed the mechanistic basis for an extensive accumulation of plasma membrane components at the cleavage furrow and the intercellular bridge. This plasma membrane accumulation is likely to decrease membrane tension at the cleavage furrow (26, 51) and generate a membrane tension gradient across the cytokinetic cell. The local changes in membrane accumulation and tension can impact several cellular processes. Most notably, decreased membrane tension at the intercellular bridge facilitates the recruitment of the abscission machinery and the completion of cell division (7, 9–13). We propose that the plasma membrane accumulation at the furrow acts as a mechanism to support the completion of cytokinesis. In addition, exocytosis and endocytosis can be triggered by high and low local membrane tensions, respectively (52–57). Based on membrane tension alone, exocytosis is favored at the cell poles and endocytosis at the cleavage furrow during ingression. This prediction agrees with experimental results in early cytokinesis (26, 58). The force-induced plasma membrane deformations revealed by our study could also result in localized regulation of mechanotransduction in cytokinesis (51, 57, 59). For example, decreased membrane tension at the intercellular bridge could influence the mechanosensitive Piezo1 channel, which localizes to the intercellular bridge and controls cytokinetic abscission (60). Furthermore, our results suggest that many of the individual membrane proteins (and lipids) that accumulate at the cleavage furrow may do so due to the general movement and folding of the plasma membrane rather than a protein (or lipid) specific accumulation mechanism. Whether similar generic membrane accumulation mechanisms also apply to other contexts where specific membrane components are observed to accumulate locally, such as cell migration, remains an interesting topic for future studies.

Exocytosis is known to take place at the intercellular bridge and is important for the completion of cytokinesis (14–20). However, our results indicate that localized exocytosis is not required for accumulating excess plasma membrane at the cytokinetic furrow. It is possible that exocytosis plays a crucial role by supplying specific components to the plasma membrane, or by secreting specific intracellular components (40). However, increasing the plasma membrane area during cytokinesis may not be required. Plasma membrane folds contain excess surface area that is estimated to be ∼50%, or more, of the apparent cell surface area (61, 62), and we observed that cells arrested to prometaphase displayed a highly folded membrane structure. As the apparent cell surface area increases ∼30% during ingression, cytokinesis could proceed in the absence of plasma membrane area addition, by relying solely on the unfolding of membrane reservoirs. Indeed, based on our surface protein labeling approach, we observed only ∼3% increase in surface components that can be attributed specifically to cytokinesis rather than cell growth. Thus, cytokinesis-specific increase in plasma membrane area is unlikely to be necessary for a successful cell division, especially when alternative mechanisms exist to decrease membrane tension at the intercellular bridge.

We find that the cortical surface experiences up to a four-fold increase in local longitudinal stretching at the cleavage furrow as the furrow constricts. This phenomenon is consistently observed across a wide range of modeling parameters, resulting in mechanical stresses on the plasma membrane that may hinder the completion of cytokinesis. Importantly, the mechanical properties of the plasma membrane can also have reciprocal effects on the actomyosin cortex (see Supplementary Note 4). The predicted longitudinal stretching at the furrow stands in contrast to previous studies from *C. elegans* embryos that suggested longitudinal compression at the division plane throughout furrow constriction (63, 64). This previous conclusion relied on measurements of actin cortex dynamics, which could misleadingly result in an apparent longitudinal cortical compression at the furrow as the cortical flows slowdown near the division plane. In contrast, our results uncouple local cell surface deformations from cortical flow effects, revealing simultaneous cell surface longitudinal stretching and slowdown of cortical components as they approach the division plane.

More broadly, our study focuses on, and is limited to, the cell intrinsic mechanical regulation of plasma membrane dynamics during cytokinesis. Given that the actomyosin contractile ring is a conserved feature among animal cells, the mechanisms that govern plasma membrane accumulation may also be accessible to all animal cells. However, the plasma membrane may not accumulate at the cleavage furrow in all cells due to differences in plasma membrane composition, actomyosin contractility, as well as cell-cell and cell-substrate interactions that further influence the behavior of the plasma membrane. The cell type dependency of our discoveries is discussed with more detail in Supplementary note 2, and limitations of our study are detailed in Supplementary Note 5.

Overall, our work indicates that the cytoskeletal forces governing cell division simultaneously accumulate plasma membrane at the cleavage furrow and the intercellular bridge. This may serve as a self-protecting mechanism against cytokinetic failures that arise from high membrane tension. Genetic and chemical manipulations that target actomyosin or membrane-cortex adhesion may therefore impact cytokinetic fidelity indirectly by regulating plasma membrane dynamics and tension.

## MATERIALS AND METHODS

### Cell lines, cell culture & chemical treatments

L1210, HeLa and BaF3 cells were cultured in RPMI (Invitrogen # 11835030) supplemented with 10% FBS (Sigma-Aldrich), 10mM HEPES, 1mM sodium pyruvate and antibiotic/antimycotic. All experiments were carried out with exponentially growing cells at a confluency of 300.000 – 600.000 cells/ml. All cell lines tested negative for mycoplasma.

For chemical treatments of live cells, we used 1:1000 dilution (v/v) of DMSO (vehicle for all chemicals, except Brefeldin A), 1:1000 dilution (v/v) of EtOH (vehicle for Brefeldin A), 50 µM (-)-Blebbistatin (Cayman Chemical, #13013) for 2 hours, 2 µM Latrunculin B (Cayman Chemical, #10010631) for 2 hours, 10 µM Brefeldin A (ThermoFisher, #00-4506-51) for 1 hour, or 50 µM CK-666 (Cayman Chemical, #29038) for 1 hour. For live cells experiments, the chemicals were present in media during imaging.

### Plasma membrane protein and lipid labeling

Plasma membrane proteins on the external side of the membrane were labeled using the LIVE/DEAD Fixable Red Dead Cell Stain kit (Invitrogen, #L23102), which labels free amine groups, using 5x supplier’s recommended concentration. The cells were washed with cold PBS and stained in cold PBS in dark at +4°C for 10 minutes. Cell washes were carried out using short and weak centrifugation (1 min at 300 g), to minimize cell shape perturbations. Staining was stopped by mixing the cells with FBS containing media, after which the cells were washed and imaged. For co-labeling with plasma membrane lipid stains, a green variant of the label was used (Invitrogen, #L23101). For end-point imaging, imaging media was at +4°C, whereas for timelapse imaging media was at +37°C. Plasma membrane lipids were labeled using CellMask Deep Red Plasma membrane stain (Invitrogen, # C10046) at 10 µg/ml concentration. Staining was carried out in media at RT for 10 min, followed by a wash using culture media.

### Biophysical modeling of cytokinesis

The cytokinesis numerical code was implemented in Fortran 90, while the analysis of the model outcomes was conducted using MATLAB. Supplementary Table 1 provides a summary of the parameter values utilized in all the numerical simulations, unless otherwise noted. A comprehensive description of the model can be found in the Supplementary Note 3.

### Data and code availability

All quantifications are shown in figures. The MATLAB code used for image analysis can be found at https://github.com/alicerlam/lam-miettinen-cell-membrane-intensity. All imaging data and codes used for biophysical model of cytokinesis are available at a reasonable request.

Please see Supplementary Materials and Methods for full experimental and image analysis details.

## ACKNOWLEDGMENTS

This work was supported by the Koch Institute Support (core) Grant P30-CA14051 from the National Cancer Institute and by the Virginia and D.K. Ludwig Fund for Cancer Research (T.P.M.), as well as by grants U54CA210190, P01CA254849 and U54CA268069 from the National Institutes of Health (R.A.M.). We thank Prof.

S. Manalis, Prof. D. J. Odde, and Prof. P. P. Provenzano for their support and mentoring, Ms. K. Suarez and Mr.

W. Wu for technical assistance, and Dr. Kiera M. Sapp for providing research materials. We also thank the Koch Institute’s Robert A. Swanson (1969) Biotechnology Center, specifically the Microscopy Core Facility and the Peterson (1957) Nanotechnology Materials Core Facility for excellent technical support. More specifically, we would like to acknowledge the contribution of David Mankus, Margaret Bisher and Abigail Lytton-Jean in obtaining the SEM images.

## Author contributions

R.A.M. developed and carried out the computational simulations of cytokinesis. A.L. developed the image analysis pipeline and quantified the microscopy data. T.P.M. conceived and supervised the study and carried out imaging experiments. R.A.M. and T.P.M. wrote the manuscript with assistance from A.L.

