## Supplementary Information for "Cell intrinsic mechanical regulation of plasma membrane accumulation at the cytokinetic furrow"

**This PDF file includes:**

- Table of Contents
- Supplementary Materials and Methods
- Supplementary Notes 1 - 5
- Table S1 (within Supplementary Note 3)
- Figures S1 to S16
- Legends for movies S1 to S5
- SI References

**Other supporting materials for this manuscript include the following:**

- Movies S1 to S5

### SUPPLEMENTARY MATERIALS AND METHODS

#### *Cell lines and genetic reporters*

The L1210 cells were obtained from ATCC (#CCL-219), the BaF3 cells were obtained from RIKEN BioResource Center, and the HeLa cells were gifted by the laboratory of Matthew Vander Heiden at Massachusetts Institute of Technology. HeLa H2B-GFP cells were gifted by Dr. Kiera M. Sapp. L1210 cells expressing the LifeAct F-actin sensor together with the Geminin-based cell cycle indicator were generated in a previous study (1). L1210 cells expressing only the cell cycle indicator were generated in (2). The cell lines were transduced with a lentiviral vector carrying H2B-GFP encoding plasmid and puromycin resistance, graciously gifted by Dr Kiera M. Sapp. L1210 and BaF3 cell transductions were carried out using spinoculation in culture media containing 8 µg/ml polybrene (3). After spinoculation, the cells were resuspended in normal culture media and grown o/n. The spinoculation procedure was repeated the next day after which selection was started using 5 µg/ml of puromycin. After a week of selection, high GFP expressing cells were sorted using FACS.

#### *Plasma membrane and F-actin labeling*

Thiol-reactive stain labeling of cell surface proteins was carried out as with the amine-reactive stain, except using Alexa Fluor 568 C5 Maleimide (Invitrogen, #A20341) at 50 µM concentration.

Plasma membrane oligosaccharides were labeled using Alexa Fluor 350 conjugate of WGA (Invitrogen, #W11263) after washing the cells with PBS. Staining concentration was 10 µg/mL, and staining was carried out in PBS in dark at +4°C for 20 minutes, followed by a was using PBS. Cells were then moved to media and imaged.

For plasma membrane cholesterol labeling, L1210 cells were washed with PBS and fixed using 4% PFA in PBS for 10 min at RT. The cells were then washed with PBS containing 5% BSA and labeled using 100 µg/ml Filipin III in PBS for 1h at RT in dark. The cells were then washed twice with PBS and imaged in PBS immediately after. For co-labeling of surface proteins and cholesterol, cells were first labeled for surface proteins, then washed 3 times with FBS containing media, and then fixed and labeled for cholesterol as detailed above. Co-labeling of surface proteins and cholesterol was carried out in H2B-GFP expressing cells, whereas co-labeling of surface proteins and membrane lipids (using CellMask Deep Red Plasma membrane stain) was carried out in wt L1210 cells.

For F-actin labeling, BaF3 cells were fixed as detailed above, and stained with 1x Phalloidin-iFluor 405 Conjugate (Cayman Chemical, #20548) for 30 min in dark at RT. After staining the cells were washed twice with PBS before imaging. L1210 cell F-actin localization was examined by expressing LifeAct, an F-actin binding protein, coupled to an RFP (1).

#### *Western blots and biochemical cell fractionation*

L1210 wt cells were first depleted of dead cells using a Dead Cell Removal kit (Miltenyi Biotec, #130-090-101) according to manufacturer's instructions. The cells were then treated with 1 µM Latrunculin B for 10 min to enhance cell fractionation efficiency between F-actin and membrane proteins. Cells were then labeled with cell surface protein label as before, after which cells were washed 2 times with full media and once with PBS. The labeled cells were then fractionated into cytoplasmic, membrane and nuclear fractions using a Cell Fractionation Kit (Cell Signaling Technology, #9038) according to manufacturer's instructions. After fractionation, the original protein labeling fluorescence from each fraction was measured using excitation at 570 nm and emission at 630 nm using a plate reader. In addition, the protein content of each fraction was measured using BCA assay. Protein content was used to normalize the fluorescence measurements.

For Western blots, the fractionated samples were mixed with XT sample buffer (Bio-Rad, #1610791), heated at 96 °C for 5 min, and run on a 4-12% Bis-Tris gel (Bio-Rad, #3450123) for 50 min at 160 V using XT MES running buffer. Precision Plus Protein WesternC standard (Bio-Rad, #161-0376) was used as a protein ladder. The proteins were transferred onto a nitrocellulose membrane at 50 V for 2 h using Tris-Glycine transfer buffer (Bio-Rad, #161-0771). A separate gel was prepared for visualizing all proteins on the gel using InstantBlue Coomassie based staining solution (Expedeon, #ISB1L). After transfer, the membrane was washed with TBST and blocked using 5% BSA in TBST for 30 min at RT. After blocking, the membrane was incubated with a single primary antibody, washed 3 times with TBST, incubated with secondary antibody, washed 3 times with TBST, and imaged, after which the process was repeated with a new primary antibody. The primary antibodies used were anti-Rab8A (D22D8) (rabbit mAb, Cell Signaling Technology, #6975), anti-GAPDH (D4C6R) (mouse mAb, Cell Signaling Technology, #97166), and anti-β-Actin (8H10D10) (mouse mAb, Cell Signaling Technology, #3700). The secondary antibodies used were conjugated to fluorescent dyes. More specifically, we used IRDye 680RD goat anti-rabbit antibody (LI-COR, #925-68071) and IRDye 800CW donkey anti-mouse antibody (LI-COR, #925-32212). All primary antibodies were diluted 1:1000 from supplier stock solutions and incubated on the membrane o/n at 4 °C. Secondary antibodies were diluted 1:2500 from supplier stock solutions and incubated on the membrane for 1 hour at RT. All antibodies were prepared in a solution made of TBST supplemented with 5% BSA, and 0.5% $NaN_3$ (w/w).

***Fluorescence microscopy***

For all labeling approaches, cells were imaged immediately after labeling. Live cell imaging was carried out in culture media containing an additional 5% of serum, while fixed cell imaging was carried out in PBS. All samples were plated on poly-L-lysine (Sigma-Aldrich, #P8920) coated glass bottom CELLVIEW dishes (Greiner Bio-One, #627975). Imaging was carried out at RT, except for timelapse imaging which was carried out at 37°C, using DeltaVision's wide-field deconvolution microscope with standard DAPI, FITC, TRITC and APC filters. L1210 and BaF3 cells were imaged using a 100x oil-immersion objective, while HeLa cells were imaged using a 60x oil-immersion objective. The immersion oil had a refractive index of 1.516 (Cargille Laboratories).

For end-point imaging, z-slices were collected with 0.25 – 0.3 µm spacing covering typically a 15 µm height. For timelapse imaging, where phototoxicity was minimized, z-slices were collected typically with a 1 µm spacing covering typically a 10 µm height. L1210 timelapse imaging was carried out every 90 sec and HeLa timelapse imaging was carried out every 60 - 150 sec. Image deconvolution was carried out using SoftWoRx 7.0.0 software.

***Image analysis***

All images were analyzed using ImageJ (version 1.53q) and MATLAB (R2023a). Dividing cells were first identified from large images with multiple cells. For each dividing cell, the z-layer with the widest cleavage furrow was cut out using ImageJ. The resulting single z-slice images were analyzed using the MATLAB analysis pipeline detailed in Figure S1. For each cell, the division plane and plasma membrane were manually defined by a freehand line profile drawn using MATLAB's Image Processing and Computer Vision toolbox. To account for the higher and variable width of the plasma membrane (i.e., folding), the images were smoothed with a 15-pixel x 15-pixel square kernel that locally averaged pixel intensities throughout the image. After smoothing, a single pixel wide line profile of the traced plasma membrane outline was recorded at ~4000 points, and the distance between each point and the division plane was calculated. These pixel intensities were plotted as a function of distance from the division plane, so that each side of the dividing cell is analyzed separately. The data was then interpolated, and the two sides of the cell are averaged to give one trace per cell. For the analysis in Fig 1B and 5E, images were manually assigned into groups based on their cell cycle stage, as identified using the H2B-GFP marker. Pixel intensities of each cell were normalized to Z-scores, using the mean and standard deviation of the cell being analyzed. In 5E, only early cytokinetic cells were quantified to avoid bias from late cytokinetic cells, which are present in DMSO treated but not drug treated samples. For Fig 1C, images were classified manually based on cell shape and DNA morphology. For the analysis of the timelapse images (Fig. 3D, 5B), the maximum pixel intensity at the cleavage furrow and the average intensity at the cell poles were extracted from the pixel intensity vs. distance plots described above. For each cell, the timelapses were manually aligned to the onset of anaphase, and data for each cell were normalized to the average intensities observed before and at the onset of anaphase.

#### *Scanning electron microscopy*

Freely proliferating L1210 H2B-GFP cells in normal culture media were placed on a round cover slip coated with poly-L-lysine. The cells were given 15 min to adhere in RT. After this, the media was removed and replaced with PBS containing 2% glutaraldehyde for fixation. The cells were fixed for 30 min at RT, then washed with PBS and twice with 100 mM sodium cacodylate buffer at 4 °C. Post fixation was carried out with 1% osmium tetroxide in the sodium cacodylate buffer for 30 min at 4 °C. The cells were then rinsed 4 times with DI H2O to remove all buffer and fixative, after which the cells were dehydrated with stepwise ethanol (EtOH) treatment at RT. Each step lasted 5 min and EtOH concentrations used were 35%, 45%, 50%, 65%, 70%, 85%, 95% and 100%. This was followed by 15 min treatment with 50% EtOH and 50% tetramethyl silane (TMS), another 15 min treatment with 20% EtOH and 80% TMS, and two final 5 min treatments with 100% TMS. After this, the TMS was removed, and the cells were left to dry at RT o/n.

Prior to imaging, the cells were sputter coated with gold for 120 s. Imaging was carried out using Zeiss Crossbeam 540 scanning electron microscope at the Peterson (1957) Nanotechnology Materials Core Facility at the Koch Institute for Integrative Cancer Research. Images were collected using a range of imaging settings, but typical imaging was carried out with 5 kV accelerating voltage, 500 pA probe current, a working distance of 8 mm, and a magnification of approximately 4000x. For all images shown in the main figures, data was collected with a resolution of 4 nm/pixel. For other images, resolution ranged from 4 to 10 nm/pixel. Images were analyzed using ImageJ (version 1.53q).

#### *Suspended microchannel resonator (SMR)*

The SMR fabrication and operation, as well as the fluorescence setup incorporation are detailed in (3–7), while details about the measurement precision can be found in (3, 4, 8, 9). In short, the SMR is a vibrating cantilever with a microfluidic channel incorporated in the cantilever. The cantilever vibration is driven by a mechanical actuator below the silicon chip within which the cantilever is incorporated, and the vibration frequency of the cantilever is detected by a piezoresistor implanted at the base of the cantilever. As cells are flown through the microfluidic channel, the resonance frequency of the cantilever changes proportionally to the buoyant mass of the cantilever. Next to the cantilever, the microfluidic channel crosses a fluorescence excitation and detection region where the geminin-GFP and cell surface amine labeling signal are recorded. During the measurements, cells were periodically flushed in the SMR from a sample vial that was kept on ice. Buoyant mass measurements were calibrated using polystyrene beads of known size.

For data analysis, each buoyant mass measurement was paired with fluorescence signals based on the time traveled between the SMR cantilever and the fluorescence detection region. If a buoyant mass measurement paired with multiple fluorescence signals, or *vice versa*, the data was excluded. Cells with excessively high amine labeling (see Fig. 2A for an example) were excluded as dead cells. Cytokinetic cells were manually gated based on cell mass and geminin-GFP signal. Then, the cytokinetic cells were matched with G2/M cells (high geminin-GFP signal) that displayed the closest buoyant mass to the cytokinetic cells (in cases of multiple matching cells, G2/M data was excluded). This was followed by a comparison of cell surface amine labeling in the equally sized G2/M and cytokinetic cells. For the calculations of % surface protein content increase in cytokinesis, the error value displayed is the compound error derived from standard error of the mean values in G2/M and cytokinesis.

***Cell proliferation rate analysis***

Cell proliferation rates were analyzed using the IncuCyte live cell imaging system by Sartorius. Cells were imaged every 3 hours for approximately 3 days and cell confluency, as analyzed the IncuCyte software, was used as a proxy for cell number to define proliferation rate. Only imaging timepoints in the exponential growth phase were used to define the cell proliferation rate (doublings per hour). The drug treatments started 30 min prior to the start of the live cell imaging.

### SUPPLEMENTARY NOTE 1: Cell surface protein labeling

Here, we assess the specificity of our cell surface protein labeling method in targeting only plasma membrane components, with a focus on verifying that the labeling is not also labeling F-actin. We first examined the surface protein labeling upon cell death and loss of plasma membrane integrity. Compromised plasma membrane integrity was apparent in dying and blebbing cells, evidenced by intracellular labeling, with an approximate 100-fold increase in signal intensity per cell (Fig. S2A). Such cells with significantly increased labeling intensity are not included in our analyses in this study.

Next, we labeled cell surface proteins in wild type L1210 cells and biochemically separated the cell membranes from the cytoplasm to examine the subcellular localization of the cell surface labeling (Fig. S2B). The purity of the cytoplasmic and membrane fractions was examined by Western blot. The cytoplasmic protein GAPDH purified near-exclusively in the cytoplasmic fraction and the membrane protein Rab8, which also localizes to the plasma membrane (10), was found near-exclusively in the membrane fraction (Fig. S2C-D). When examining the presence of the surface label, we found that the membrane fraction displayed ~33-fold higher labeling than the cytoplasmic fraction (Fig. S2E). Thus, our amine-reactive cell surface protein labeling approach is labeling predominantly proteins on the plasma membrane. Importantly, β-Actin was found near-exclusively in the cytoplasmic fraction (Fig. S2D), indicating that the surface protein labeling is also not penetrating the plasma membrane and labeling the actin cortex.

We then treated L1210 cells with 2 μM Latrunculin B for 2 hours to depolymerize F-actin, after which we labeled cell surface proteins. If the cell surface protein labeling also targets F-actin, we would expect a decrease in the total labeling intensity following Latrunculin B pre-treatment. However, Latrunculin B pre-treatment did not alter the total labeling intensity when compared to DMSO treatment, as analyzed using flow cytometry (Fig. S2F). We also considered the situation where cells were first surface-labeled and then treated with Latrunculin B. If the cell surface protein labeling also labels F-actin, Latrunculin B treatment should result in increased cell internal labeling as the labeled actin molecules dissociate from the actin cortex. However, when examining L1210 cell internal labeling with microscopy, we observed a decrease in the cell cytosolic signal following an exposure to Latrunculin B as opposed to DMSO treatment (Fig. S2G).

In BaF3 cells, approximately 50% of cells displayed cell surface protein accumulation at the cytokinetic furrow (Fig. 1C-D), and approximately 60% of cells displayed cell surface lipid accumulation at the cytokinetic furrow (Fig. S4A,B,D,E). In contrast, almost all (~95%) of the BaF3 cells displayed F-actin accumulation at the cytokinetic furrow (Fig. S4C). This discrepancy between the accumulation of plasma membrane components and F-actin at the cytokinetic furrow (Fig. S4E) indicates that our plasma membrane labeling approaches are independent of F-actin.

Finally, we verified that our results on global accumulation of plasma membrane proteins at the cleavage furrow are not specific to our approach for labeling cell surface proteins. Instead of the amine-reactive label, we used a thiol-reactive, cell impermeable, fluorescent label. This labeling revealed similar plasma membrane protein accumulation at the cytokinetic furrow of L1210 cells as observed with the amine-reactive label (Fig. S2H).

Together, these results show that our cell surface protein labeling is not binding to F-actin or other cytoplasmic components, thereby allowing us to interpret cell surface protein labeling results as plasma membrane protein movements that are separate from F-actin movements on the cell cortex.

**SUPPLEMENTARY NOTE 2: Mechanical explanations for cell type differences in cytokinetic plasma membrane dynamics**

Our results revealed that all L1210 cells display plasma membrane accumulation at the cytokinetic furrow but only approximately half of BaF3 and HeLa cells display this phenotype when carrying out end-point imaging of cytokinetic cells (Fig. 1C). More broadly, previous literature suggests that plasma membrane does not accumulate at the cytokinetic furrow in all cell types (11). Such phenotype heterogeneity is not uncommon for membrane dynamics. For example, intracellular vesicle trafficking to the intracellular bridge has been reported to take place in ~60% of BSC1 cells (12). Moreover, in the case of HeLa cells, our timelapse imaging (Fig. S6) results revealed that most HeLa cells (13 out of 15 cells) accumulated plasma membrane in the cytokinetic furrow, but this effect did not persist throughout cytokinesis. While mechanical stresses on the plasma membrane that cell adhesion generates can explain the weakened effect size in HeLa cells, it does not explain why all BaF3 cells, which grow in suspension, did not display the membrane accumulation at the cytokinetic furrow. We therefore sought to identify conditions where plasma membrane accumulation at the cytokinetic furrow would be diminished.

Different cell types are known to display a wide range of mechanical properties of their actin cortex and plasma membrane (13, 14). Based on our results detailed in the main text (Fig. 5A), low membrane accumulation would require both strong attenuation of membrane flows and minimization of surface area constriction. The strength of directed membrane flows towards the furrow is mainly regulated by membrane fluidity. Consequently, a high membrane lipid-protein drag coefficient $\gamma^{\mathrm{m}}_{\mathrm{drag}}$ would be sufficient to effectively attenuate membrane flows and suppress in-plane transport-mediated membrane accumulation at the furrow. In addition to altered membrane flows, the extent of localized surface area compression at the furrow can also change. The local surface area compression is largely governed by the tension exerted by the contractile ring (Fig. 4F). Increasing contractile ring tension results in enhanced longitudinal stretching that can counteract the radial area compression, resulting in minimal cell surface area changes at late stages of furrow ingression (Fig. S10B-C). Accordingly, minimal membrane content at the furrow is observed for high cortical contractility ($\sigma^{\mathrm{c,act}}_{\mathrm{max}} > 1100\ \mathrm{pN} \cdot \mu\mathrm{m}^{-1}$) and low membrane-cortex adhesion ($\gamma^{\mathrm{m-c}}_{\mathrm{drag}} < 30\ \mathrm{pN} \cdot \mathrm{s} \cdot \mu\mathrm{m}^{-3}$) (Fig. S10D-E). Both high cortical contractility and low membrane-cortex adhesion are necessary to observe minimal accumulation of membrane content at the furrow. Overall, these results suggest that the cell-to-cell differences we observe between the three cell lines studied in this work could arise simply from differences in cortical contractility, membrane-cortex linker levels and/or membrane fluidity. Notably, cells can regulate these mechanical properties and, consequently, the capacity for plasma membrane accumulation at the cytokinetic furrow may exist also in cells that do not display the membrane accumulation.

**SUPPLEMENTARY NOTE 3: Force balance model of cytokinesis**

**Theoretical model of cytokinesis**

We extend previous theoretical work (15, 16) and develop a biophysical model of cytokinesis. The framework treats the actomyosin cortex as a continuous surface rather than resolving individual cortical components. This simplification provides a more computationally efficient way to predict the behavior of the cell serving as a bridge between microscopic and macroscopic scales. Furthermore, this modeling approach permits cell-specific parameterization using available data, enabling accurate predictions of cell behavior and large-scale cell deformations that occur during cytokinesis. We model the cortex as a thin actomyosin viscous fluid film that generates stresses. The constitutive equation for the cortical stress can be written as the sum of a viscous contribution and an active contribution: $\sigma_{ij}^{c} = \sigma_{ij}^{c,visc} + \sigma_{ij}^{c,act}$, where i and j are tensor indices. Viscous shear stresses can arise from in-plane velocity gradients or by surface deformation. The constitutive equation for the viscous stress $\sigma_{ij}^{c,visc}$ reads:

$$\sigma_{ij}^{c,visc} = \eta^{c}\left(\nabla_i v_j^{c} + \nabla_j v_i^{c}\right) + 2\eta^{c} v_n C_{ij}. \tag{S1}$$

Here, $\eta^{c}$ is the cortex viscosity, $\nabla_i$ is the covariant derivative, $v_j^{c}$ is the in-plane cortical flow velocity, $v_n$ is the surface normal velocity that captures cell surface deformations, and $C_{ij} = \mathbf{e_j} \cdot \partial_i \mathbf{n}$ is the second fundamental form of the cell surface, or curvature tensor. Here, $\mathbf{e_j}$ are the basis vectors in the tangent space defined by the surface and $\mathbf{n}$ is the outward unit vector normal to the cell surface (17, 18). The constitutive equation for the cortical active stress $\sigma_{ij}^{c,act}$ reads:

$$\sigma_{ij}^{c,act} = \sigma_{max}^{c,act} \operatorname{erf}\left(\frac{\rho^{act}}{\rho_s^{act}}\right) \operatorname{erf}\left(\frac{\rho^{myo}}{\rho_s^{myo}}\right) g_{ij}. \tag{S2}$$

The parameter $\sigma_{max}^{c,act}$ defines the maximum cortical tension that the cortex can generate. We assume that cortical tension is a function of the local actin density $\rho^{act}$, and myosin density $\rho^{myo}$, with saturation behavior. This means that the tension in the network becomes less sensitive to changes in actin and myosin density as their densities approach their saturation values, $\rho_s^{act}$ and $\rho_s^{myo}$, respectively. Actin and myosin densities represent the number of actin units and myosin molecules per unit surface area, respectively. The tensor $g_{ij}$ introduced in Eq. (S2) represents the first fundamental form of the surface, also called metric tensor, which can be defined as $g_{ij} = \mathbf{e_i} \cdot \mathbf{e_j}$. Normal force balance on the cortex reads:

$$-\sigma^{c,ij} C_{ij} = \Delta p, \tag{S3}$$

where $\Delta p$ is a pressure term that quantifies the pressure difference between the intracellular and extracellular spaces. Mathematically, it behaves as a Lagrange multiplier, a scalar quantity that adjusts its value to ensure the cell volume always remains constant. The tangential force balance on the cortex can be written as:

$$\nabla_i \sigma^{c,ij} - \gamma_{drag}^{m-c}\left(v^{c,j} - v^{m,j}\right) = 0. \tag{S4}$$

This equation describes how active forces drive cortical flows, which are resisted by cortex viscous forces and drag forces originating from the cortex sliding along the plasma membrane. Drag forces are mediated by membrane-cortex linkers and proportional to the difference between the in-plane cortical flow velocity $v^{c,j}$ and the in-plane membrane lipid flow velocity $v^{m,j}$, with constant of proportionality represented by an effective membrane-cortex drag coefficient $\gamma_{drag}^{m-c}$. We assume that the parameter $\gamma_{drag}^{m-c}$ maintains a constant value over time and across the entire cell. Solution of the force balance equations allows us to compute cortical flows and cell shape deformations, given by the cortical flow velocity $v^{c,i}$ and deformation velocity $v_n$, respectively. We treat the cell surface as a Lagrangian structure, described by the shape position vector $\mathbf{X}$. Cell surface deformations are given by:

$$\frac{d\mathbf{X}}{dt} = v_n \mathbf{n} \tag{S5}$$

The spatial and temporal changes in the density of cortical actin is described by the following mass conservation equation:

$$\frac{\partial \rho^{act}}{\partial t} = k_{on}^{act} \rho_{free}^{act} - k_{off}^{act} \rho^{act} - \nabla_i\left(\rho^{act} v^{c,i}\right) - C_i^i \rho^{act} v_n + D^{act} \nabla_i \nabla^i \rho^{act}. \tag{S6}$$

The term on the left-hand-side (LHS) of Eq. (S6) represents the rate-of-change of actin density. The first two terms on the right-hand-side (RHS) capture simple turnover kinetics, where $k_{on}^{act}$ and $k_{off}^{act}$ are the actin association and dissociation rate constants, respectively. The concentration of actin in the cytoplasm has been denoted as $\rho_{free}^{act}$. Notice that the total number of actin units in the cell $N_{tot}^{act}$ is equal to the sum of the free pool of actin in the cytoplasm and the number of cortical actin units. This can be expressed mathematically as $N_{tot}^{act} = \rho_{free}^{act} V_{cell} + \int dS \rho^{act}$. Here, $V_{cell}$ is the cell volume and the surface integral over the whole cell is used to calculate the total number of actin units in the cortex. The third term in Eq. (S6) represents the transport of actin powered by in-plane cortical flows, the fourth term represents the rate-of-change of actin caused by local cell surface area changes, and the last term captures actin diffusion, with diffusion constant $D^{act}$. Similarly, we write down the mass conservation equation for myosin as:

$$\frac{\partial \rho^{myo}}{\partial t} = k_{on}^{myo} \rho_{free}^{myo} - k_{off}^{myo} \rho^{myo} - \nabla_i\left(\rho^{myo} v^{c,i}\right) - C_i^i \rho^{myo} v_n + D^{myo} \nabla_i \nabla^i \rho^{myo}, \tag{S7}$$

where we account for myosin association and dissociation kinetics, in-plane myosin transport, accumulation or dilution of myosin due to local cell surface area changes and in-plane diffusive processes. $k_{on}^{myo}$ and $k_{off}^{myo}$ are the myosin association and dissociation rate constants, respectively, $\rho_{free}^{myo}$ is the concentration of myosin in the cytoplasm, and $D^{myo}$ is the myosin diffusion constant. The overall count of myosin molecules within the cell comprises both the cytoplasmic and cortical pools: $N_{tot}^{myo} = \rho_{free}^{myo} V_{cell} + \int dS \rho^{myo}$.

Next, we proceed to model plasma membrane mechanics and dynamics. The membrane contains lipids and membrane proteins. We treat it as a continuous surface, as opposed to individually tracking individual lipid molecules, and assume that it behaves as a two-dimensional viscous fluid layer with elastic properties and subjected to drag forces. At the microscopic level, the plasma membrane contains folded regions (Fig. 2E, Main Text), and other membrane reservoirs. To quantify local plasma membrane amounts, we define the apparent cell surface contour as determined by the cortex configuration, and we introduce a projected lipid density $\rho^m$, a variable that represents the total number of lipids projected onto the cell surface per unit of apparent cell surface area.

We assume that the constitutive equation for the membrane stress takes the following form:

$$\sigma_{ij}^m = \eta^m\left(\nabla_i v_j^m + \nabla_j v_i^m\right) + 2\eta^m v_n C_{ij} - E^m\left(\frac{\rho^m - \rho_0^m}{\rho_0^m}\right) g_{ij}. \tag{S8}$$

Here, $\eta^m$ is the membrane viscosity and $v_j^m$ is the net in-plane lipid flow velocity. The first two terms on the RHS of Eq. (S8) represent viscous stresses that arise from in-plane velocity gradients and cell surface deformations. The third term represents membrane tension, which assumes a linear stress-strain relation (19). Membrane tension tends to reduce lipid density inhomogeneities by smoothing folded membrane structures. We have denoted the effective membrane elastic modulus coefficient as $E^m$ and the unperturbed projected lipid density as $\rho_0^m$, which is the projected density of lipids in the absence of net flow of lipids and cell surface deformations. According to the original fluid mosaic model, plasma membrane constituents freely diffuse within the plane of the membrane (20). However, the diffusion coefficients associated to many transmembrane proteins are nearly two orders of magnitude smaller than those found in artificially reconstituted membranes and liposomal membranes (21, 22). Therefore, in-plane lipid flows through the crowded membrane are powered by cortical flows and hindered by nearly immobile embedded proteins (23). Tangential force balance on the membrane thus reads:

$$\nabla_i \sigma^{m,ij} - \gamma_{drag}^{m-c}\left(v^{m,j} - v^{c,j}\right) - \gamma_{drag}^{m} v^{m,j} = 0. \tag{S9}$$

The first term in Eq. (S9) represents viscous forces and membrane tension forces, the second term captures membrane lipid-cytoskeletal drag forces, and the third term represents the drag forces experienced by lipids as the encounter the immobile embedded proteins, where $\gamma_{drag}^m$ is the plasma membrane lipid-protein drag coefficient. The rate-of-change of the projected two-dimensional lipid density obeys the following transport equation:

$$\frac{\partial \rho^m}{\partial t} = -\nabla_i\left(\rho^m v^{m,i}\right) - C_i^i \rho^m v_n. \tag{S10}$$

The first two terms on the RHS represent the local change of projected lipid amounts due to in-plane lipid flows and local cell surface area changes, respectively. A local cell surface area compression induces membrane folding resulting in lipid accumulation and an increase in the projected lipid density, whereas a local cell surface area expansion induces membrane unfolding resulting in lipid dilution and a decrease in the projected lipid density. These membrane folding and lipid accumulation predictions are independent of endo- and exocytosis, which are not included in the model.

**Surface parameterization and numerical approach**

We adopt a local coordinate system $\{s, \theta\}$ to describe the cell surface and create a map that assigns each coordinate pair $\{s, \theta\}$ to a point in three-dimensional space, which we describe by the shape position vector $\mathbf{X}(s, \theta, t)$, where s is the arc length parameter, $\theta$ is the azimuthal angle and t denotes time (Fig. S14). We assume that the cell surface remains axisymmetric and use the following arc-length parameterization (15, 24):

$$\mathbf{X}(s, \theta, t) = r(s, t)\bar{\mathbf{e}}_{\mathbf{r}}(\theta) + z(s, t)\bar{\mathbf{e}}_{\mathbf{z}}, \tag{S11}$$

where $r(s, t)$ and $z(s, t)$ are the radial and axial coordinates of the cell surface, respectively, and $\bar{\mathbf{e}}_{\mathbf{r}}$ and $\bar{\mathbf{e}}_{\mathbf{z}}$ are the normalized cylindrical coordinates basis vectors. Here, $s \in [0, L(t)]$ and $\theta \in [0,2\pi]$, where $L(t)$ is the cell pole-to-cell pole distance following the cell surface curve. In cartesian coordinates, the cell surface vector reads:

$$\mathbf{X}(s, \theta, t) = \begin{pmatrix} r(s,t)\cos\theta \\ r(s,t)\sin\theta \\ z(s,t) \end{pmatrix} \tag{S12}$$

To simplify our notation, we introduce the variable $\alpha(s, t)$ which is the angle of the surface curve relative to the radial axis. Notice that $\cos\alpha = \partial_s r(s, t)$, and $\sin\alpha = \partial_s z(s, t)$. The basis vectors that comprise the coordinate system on every point of the surface can be written as:

$$\mathbf{e}_{\mathbf{s}} = \partial_s \mathbf{X} = \begin{pmatrix} \cos\alpha\cos\theta \\ \cos\alpha\sin\theta \\ \sin\alpha \end{pmatrix}, \qquad \mathbf{e}_{\boldsymbol{\theta}} = \partial_\theta \mathbf{X} = \begin{pmatrix} -r(s,t)\sin\theta \\ r(s,t)\cos\theta \\ 0 \end{pmatrix} \tag{S13 − S14}$$

The associated surface normal unit vector is given by:

$$\mathbf{n} = \mathbf{e}_{\boldsymbol{\theta}} \times \mathbf{e}_{\mathbf{s}} / |\mathbf{e}_{\boldsymbol{\theta}} \times \mathbf{e}_{\mathbf{s}}| = \begin{pmatrix} \sin\alpha\cos\theta \\ \sin\alpha\sin\theta \\ -\cos\alpha \end{pmatrix} \tag{S15}$$

We proceed to calculate some geometric objects that we will use to write the model equations in axisymmetric form using the surface parameterization introduced in Eq. (10). The first and second fundamental forms of the surface are given by:

$$\begin{pmatrix} g_{\theta\theta} & g_{\theta s} \\ g_{s\theta} & g_{ss} \end{pmatrix} = \begin{pmatrix} r^2 & 0 \\ 0 & 1 \end{pmatrix} \tag{S16}$$

$$\begin{pmatrix} C_{\theta\theta} & C_{\theta s} \\ C_{s\theta} & C_{ss} \end{pmatrix} = \begin{pmatrix} \dfrac{\sin\alpha}{r} & 0 \\ 0 & \dfrac{\partial\alpha}{\partial s} \end{pmatrix} \tag{S17}$$

Due to the spatial dependence of the coordinate system on curvilinear manifolds, derivatives of vectors and tensors must be corrected by some terms that involve the so-called Christoffel symbols of the first kind and second kind, typically denoted by the symbols $\Gamma_{ijk}$ and $\Gamma_{ij}^{k}$, respectively. They are defined as $\Gamma_{ijk} = \frac{1}{2}\left[\partial_i g_{jk} + \partial_j g_{ki} + \partial_k g_{ij}\right]$ and $\Gamma_{ij}^{k} = g^{kl}\Gamma_{ijl}$. In our axisymmetric analysis, we only make use of the Christoffel symbol $\Gamma_{\theta s}^{\theta}$, whose expression is given by $\Gamma_{\theta s}^{\theta} = \cos\alpha / r$.

Because the arc-length domain changes over time, it is convenient to introduce a Eulerian surface parameterization:

$$\mathbf{X_e}(u,\theta,t) = r(u,t)\bar{\mathbf{e}}_{\mathbf{r}}(\theta) + z(u,t)\bar{\mathbf{e}}_{\mathbf{z}}, \qquad \text{(S18)}$$

where u is defined on the fixed interval $u \in [0, L(0)]$ and represents the arc-length parameter in the undeformed configuration. The arc-length parameters s and u are related by the following time-dependent coordinate transformation $\varepsilon_s(u,t)$, defined as:

$$s(u,t) = \int_0^u \varepsilon_s(u',t)du'. \qquad \text{(S19)}$$

The variable $\varepsilon_s(u,t)$ quantifies the straining in the direction of the axis of revolution of a small surface element defined by the fixed Eulerian coordinate u over time t. If $\varepsilon_s < 1$, the surface element undergoes longitudinal compression, whereas if $\varepsilon_s > 1$, the surface element undergoes longitudinal stretching. Its temporal evolution is given by (15):

$$\frac{\partial \varepsilon_s}{\partial t} = \varepsilon_s C_s^s v_n, \qquad \varepsilon_s(u,0) = 1. \qquad \text{(S20)}$$

To quantify local changes in cell surface area, we introduce the cell surface area factor $A(u,t)$, which quantifies the degree of deformation of a small cell surface element defined by the coordinate u over time t. The temporal evolution of A reads:

$$\frac{\partial A}{\partial t} = A C_i^i v_n, \qquad A(u,0) = 1. \qquad \text{(S21)}$$

If $A < 1$, the surface element undergoes compression, whereas if $A > 1$, the surface element undergoes expansion. As an example, $A(u,t) = 2$ indicates that the local cell surface area at a cell location given by u has increased two-fold during time t, whereas $A(u,t) = 0.5$ indicates that the local cell surface area decreased two-fold during time t.

Next, we rewrite the model equations (S1−S10) on the axisymmetric cell surface by using the Eulerian surface parameterization introduced in Eq. (S18). Tangential and normal force balance on the cortex read:

$$\frac{2\eta^c}{{\varepsilon_s}^2}\frac{\partial^2 v^c}{\partial u^2} + 2\eta^c\left(\frac{\Gamma_{\theta s}^{\theta}}{\varepsilon_s} - \frac{1}{{\varepsilon_s}^3}\frac{\partial \varepsilon_s}{\partial u}\right)\frac{\partial v^c}{\partial u} - \left[2\eta^c {\Gamma_{\theta s}^{\theta}}^2 + \gamma_{drag}^{m-c}\right]v^c + \frac{2\eta^c}{\varepsilon_s} C_s^s \frac{\partial v_n}{\partial u} +$$

$$+2\eta^{c}\left[\frac{1}{\varepsilon_s}\frac{\partial C_s^s}{\partial u}+\Gamma_{\theta s}^{\theta}\left(C_s^s-C_\theta^\theta\right)\right]v_n = -\frac{1}{\varepsilon_s}\frac{\partial \sigma^{c,act}}{\partial u}-\gamma_{drag}^{m-c}\,v^m \tag{S22}$$

$$\frac{2\eta^c C_s^s}{\varepsilon_s}\frac{\partial v^c}{\partial u}+2\eta^c C_\theta^\theta \Gamma_{\theta s}^\theta v^c+2\eta^c\left({C_s^s}^2+{C_\theta^\theta}^2\right)v_n=\Delta p-\sigma^{c,act}\left(C_s^s+C_\theta^\theta\right) \tag{S23}$$

Similarly, the tangential force balance equation on the plasma membrane reads:

$$\frac{2\eta^m}{{\varepsilon_s}^2}\frac{\partial^2 v^m}{\partial u^2}+2\eta^m\left(\frac{\Gamma_{\theta s}^\theta}{\varepsilon_s}-\frac{1}{{\varepsilon_s}^3}\frac{\partial \varepsilon_s}{\partial u}\right)\frac{\partial v^m}{\partial u}-\left[2\eta^m{\Gamma_{\theta s}^\theta}^2+\gamma_{drag}^{m-c}+\gamma_{drag}^{m}\right]v^m=$$
$$=-\frac{2\eta^m}{\varepsilon_s}\frac{\partial (v_n C_s^s)}{\partial u}-\;2\eta^m\Gamma_{\theta s}^\theta\left(C_s^s-C_\theta^\theta\right)v_n+\frac{E^m}{\rho_0^m\varepsilon_s}\frac{\partial \rho^m}{\partial u}-\gamma_{drag}^{m-c}\,v^{c,j} \tag{S24}$$

The time-evolution of geometric surface properties is given by the following differential equations:

$$\frac{\partial r}{\partial t}=v_n r\,C_\theta^\theta \tag{S25}$$

$$\frac{\partial C_s^s}{\partial t}=-{C_s^s}^2 v_n-\frac{1}{\varepsilon_s}\frac{\partial}{\partial u}\left(\frac{1}{\varepsilon_s}\frac{\partial v_n}{\partial u}\right) \tag{S26}$$

$$\frac{\partial C_\theta^\theta}{\partial t}=-{C_\theta^\theta}^2 v_n-\frac{\Gamma_{\theta s}^\theta}{\varepsilon_s}\frac{\partial v_n}{\partial u} \tag{S27}$$

$$\frac{\partial \Gamma_{\theta s}^\theta}{\partial t}=C_\theta^\theta\left(\frac{1}{\varepsilon_s}\frac{\partial v_n}{\partial u}-\;\Gamma_{\theta s}^\theta\,v_n\right) \tag{S28}$$

$$\frac{\partial \varepsilon_s}{\partial t}=\varepsilon_s C_s^s v_n \tag{S29}$$

Time-evolution of actin, myosin and plasma membrane lipids is given by the following mass conservation equations:

$$\frac{\partial \rho^{act}}{\partial t}=k_{on}^{act}\rho_{free}^{act}-k_{off}^{act}\rho^{act}-\frac{1}{\varepsilon_s}\frac{\partial}{\partial u}\left(\rho^{act}v^c\right)-\Gamma_{\theta s}^\theta\rho^{act}v^c-\left(C_s^s+C_\theta^\theta\right)\rho^{act}v_n+$$
$$+D^{act}\left[\frac{1}{{\varepsilon_s}^2}\frac{\partial^2\rho^{act}}{\partial u^2}+\frac{1}{\varepsilon_s}\left(\Gamma_{\theta s}^\theta-\frac{1}{{\varepsilon_s}^2}\right)\frac{\partial \varepsilon_s}{\partial u}\frac{\partial \rho^{act}}{\partial u}\right] \tag{S30}$$

$$\frac{\partial \rho^{myo}}{\partial t}=k_{on}^{myo}\rho_{free}^{myo}-k_{off}^{myo}\rho^{myo}-\frac{1}{\varepsilon_s}\frac{\partial}{\partial u}\left(\rho^{myo}v^c\right)-\Gamma_{\theta s}^\theta\rho^{myo}v^c-\left(C_s^s+C_\theta^\theta\right)\rho^{myo}v_n+$$
$$+D^{myo}\left[\frac{1}{{\varepsilon_s}^2}\frac{\partial^2\rho^{myo}}{\partial u^2}+\frac{1}{\varepsilon_s}\left(\Gamma_{\theta s}^\theta-\frac{1}{{\varepsilon_s}^2}\right)\frac{\partial \varepsilon_s}{\partial u}\frac{\partial \rho^{myo}}{\partial u}\right] \tag{S31}$$

$$\frac{\partial \rho^{m}}{\partial t} = -\frac{1}{\varepsilon_s}\frac{\partial}{\partial u}(\rho^{m}v^{m}) - \Gamma_{\theta s}^{\theta}\rho^{m}v^{m} - \left(C_s^{s} + C_\theta^{\theta}\right)\rho^{m}v_n. \tag{S32}$$

We compute the cytoplasmic actin and myosin concentrations inside the cell at each timestep as:

$$\rho_{free}^{myo} = \frac{1}{V_{cell}}\left(N_{tot}^{myo} - 2\pi\int_0^{L(0)} \rho^{myo} r\varepsilon_s du\right) \tag{S33}$$

$$\rho_{free}^{act} = \frac{1}{V_{cell}}\left(N_{tot}^{act} - 2\pi\int_0^{L(0)} \rho^{act} r\varepsilon_s du\right) \tag{S34}$$

We used $N_u$ equally spaced points to discretize the meridional cell centerline, in the fixed arc-length domain $u \in [0, \pi R_{cell}]$. The cell is initially spherical with cell radius $R_{cell}$. To mimic an enhancement of contractility at the equatorial region, we use the following actin and myosin densities as initial conditions:

$$\rho^{myo}(s_i, 0) = \rho_0^{myo} + 0.01e^{-\left[\frac{R_{cell}\sin\alpha_i}{0.3}\right]^2}, \quad \rho^{act}(s_i, 0) = \rho_0^{act} + 0.01e^{-\left[\frac{R_{cell}\sin\alpha_i}{0.3}\right]^2}, \quad (S35 - S36)$$

where $\alpha_i = \alpha(s_i, 0)$, $s_i$ is the arc-length associated to the collocation points of the initial spherical surface. We discretized spatial derivatives using second-order center finite differences and performed time-marching of model equations $(S22 - S34)$ by using an explicit first-order accurate Euler-scheme. The numerical code was implemented in Fortran.

**Estimation of model parameters**

The different model parameters are estimated based on literature background, as detailed in Table S1. We also obtained experimental support for our choice of model parameters. We treated L1210 cells with CK-666, an Arp2/3 inhibitor which increases cortical contractility and myosin penetration in the actin cortex (14, 25, 26). Our model predicts that increasing cortical contractility will not influence plasma membrane accumulation at the cytokinetic furrow in cells with high membrane fluidity ($\gamma_{drag}^{m} \sim 100 pN \cdot s \cdot \mu m^{-3}$) (Fig. 5A) but will do so in cells with low membrane fluidity ($\gamma_{drag}^{m} \sim 4000 pN \cdot s \cdot \mu m^{-3}$) (Supplementary Note 2, Fig. S10D). Increasing cortical contractility did not alter membrane accumulation at the cytokinetic furrow despite impacting cell proliferation (Fig. S11), supporting our use of low membrane lipid-protein drag coefficients.

**Table S1. Model parameters**

| Symbol | Description | Estimated values | | Legend/References |
|---|---|---|---|---|
| $N_{tot}^{act}$ | Total number of actin units in the cell | $1.14 \times 10^{7}$ | | **A** |
| $k_{on}^{act}$ | Actin association rate constant | $838.64 \mu M^{-1} \cdot s^{-1} \cdot \mu m^{-2}$ | | **B** |
| $k_{off}^{act}$ | Actin dissociation rate constant | $0.3 s^{-1}$ | | (27) |

| | | | | |
|---|---|---|---|---|
| $D^{act}$ | Actin diffusion constant | $1\mu m^2 \cdot s^{-1}$ | | **C** |
| $N_{tot}^{myo}$ | Total number of myosin molecules in the cell | $10^6$ | | **D** |
| $k_{on}^{myo}$ | Myosin association rate constant | $36.36\mu M^{-1} \cdot s^{-1} \cdot \mu m^{-2}$ | | **E** |
| $k_{off}^{myo}$ | Myosin dissociation rate constant | $0.07s^{-1}$ | | (27, 28) |
| $D^{myo}$ | Myosin diffusion constant | $1\mu m^2 \cdot s^{-1}$ | | **C** |
| $\eta^c$ | Cortex viscosity | $50pN \cdot s \cdot \mu m^{-1}$ | | **F** |
| $\sigma_{max}^{c,act}$ | Maximum actomyosin cortical contractility | $800pN \cdot \mu m^{-1}$ | | **G** |
| $\rho_s^{act}$ | Actin saturation projected density | $2.5\rho_0^{act} = 3.075 \times 10^4\mu m^{-2}$ | | **C** |
| $\rho_s^{myo}$ | Myosin saturation projected density | $2.5\rho_0^{myo} = 909\mu m^{-2}$ | | **C** |
| $\gamma_{drag}^{m-c}$ | Plasma membrane lipid-cortex drag coefficient | $40pN \cdot s \cdot \mu m^{-3}$ | | **F** |
| $\gamma_{drag}^{m}$ | Plasma membrane lipid-protein drag coefficient | $100pN \cdot s \cdot \mu m^{-3}$ | | **F** |
| $\eta^m$ | Plasma membrane viscosity | $3 \times 10^{-3}pN \cdot s \cdot \mu m^{-1}$ | | (23) |
| $E^m$ | Effective plasma membrane area expansion modulus | $40pN \cdot \mu m^{-1}$ | | (23, 29, 30) |
| $\rho_0^m$ | Unperturbed projected lipid density | $5 \times 10^6\mu m^{-2}$ | | (31) |
| $R_{cell}$ | Initial cell radius | $7\mu m$ | | **H** |
| $N_u$ | Number of discretization points | 140 | | **C** |

Cortex parameters are shown with no shade, plasma membrane parameters are shown with light gray shade, and other model parameters are shown with dark gray shade.

**A.** The total actin concentration in the cell has been recently measured in *Saccharomyces cerevisiae*: $\rho_{tot_0}^{act} = 13.2\mu M$ (32). The total number of actin units in the cell is then $N_{tot}^{act} = 1.14 \times 10^7$.

**B**. We estimate the steady-state free actin concentration in the cell as that of *Saccharomyces cerevisiae*: $\rho_{free_0}^{act} = 4.4\mu M$. The steady-state number of free actin units in the cytoplasm is then $N_{cyto_0}^{act} = \rho_{free_0}^{act} V_{cell} = 3.82 \times 10^6$. We compute the cortical actin density at steady-state conditions as $\rho_0^{act} = (N_{tot}^{act} - N_{cyto_0}^{act})/A_{cell} = 1.23 \times 10^4\mu m^{-2}$. We can estimate the rate constant at which actin units are added to the cortex from a kinetic balance of actin in the cortex: $k_{on}^{act} = \rho_0^{act} k_{off}^{act} / \rho_{free_0}^{act} = 838.64 \mu M^{-1} \cdot s^{-1} \cdot \mu m^{-2}$.

**C.** Arbitrarily chosen.

**D.** We assume that the density of myosin molecules in the cortex is that of the fission yeast cytokinetic ring (33), $\rho_0^{myo} = 500/11 \mu m \times 0.125\, \mu m = 363.6 \mu m^{-2}$. The cytoplasmic concentration of the different nonmuscle myosin II isoforms has been measured in HeLa cells and a few pancreatic cancer cell lines (34). Based on these measurements, the total free myosin concentration falls within the range $\rho_{myo}^{free} \sim [0.6 - 0.8]\ \mu M$. For our steady-state estimations, we use $\rho_{free\ 0}^{myo} = 0.7 \mu M$. We can now estimate the total number of myosin molecules in the cell as the sum of the free pool of myosin in the cytoplasm and the number of myosin molecules in the cortex: $N_{tot}^{myo} = \rho_{free\ 0}^{myo} V_{cell} + \rho_0^{myo} A_{cell} = 8.3 \times 10^5$. We use $N_{tot}^{myo} = 10^6$.

**E.** We estimate the myosin association rate constant from a kinetic balance of myosin in the cortex $k_{on}^{myo} = \rho_0^{myo} k_{off}^{myo} / \rho_{free\ 0}^{myo} = 36.36 \mu M^{-1} \cdot s^{-1} \cdot \mu m^{-2}$.

**F.** Screened parameter.

**G.** Reported values of cortical tension lie within the range $\sigma^c \approx [400 - 1600]\ pN \cdot \mu m^{-1}$ (14, 35, 36). Unless otherwise noted, we use $\sigma_{max}^{c,act} = 800 pN \cdot \mu m^{-1}$.

**H.** Most L1210 cells display a volume of $V_{cell} = [1400 - 1600]$ fL at the onset of cytokinesis, when excluding volume increase driven by the mitotic cell swelling (37). This volume range corresponds to a cell radius of $R_{cell} = [6.93 - 7.26] \mu m$. We set the cell radius $R_{cell}$ to 7μm in all our simulations unless otherwise noted.

### SUPPLEMENTARY NOTE 4: Additional insights from the model

#### Cortical tension anisotropy and plasma membrane accumulation at the cytokinetic furrow.

Previous theoretical results (38) and this work show that furrow ingression does not require the generation of anisotropic cortical tension in the division plane. Rather, a gradient of cortical tension from the cell poles to the cell's equator is sufficient to drive furrow ingression. However, actin filaments exhibit significant circumferential alignment in the division plane during cytokinesis. Cortical compression by flow (39), myosin-based search and capture (40), or pre-alignment by compression followed by formin-mediated growth of highly oriented actin filaments (41) have been proposed as potential mechanisms responsible for equatorial filament alignment.

Irrespective of the dominant mechanism of equatorial filament alignment, we aimed to explore the impact of increased circumferential tension at the division plane on local cell surface area changes and accumulation of plasma membrane components. To mimic actomyosin ring assembly and increased local filament growth along the circumferential direction, we increased the circumferential actomyosin tension in a small region around the equator. The modified circumferential cortical active stress reads:

$$\sigma_{\theta\theta}^{c,act} = \sigma_{ss}^{c,act}\left[1 + \chi_{ani} e^{-0.5\left([s-s_{eq}]/h_{ani}\right)^2}\right] \qquad \text{(S37)}$$

where $\sigma_{ss}^{c,act}$ is the axial cortical active stress given by Eq. (S2), $\chi_{ani}$ is the anisotropic parameter strength, $h_{ani}$ is the characteristic anisotropic length and $s_{eq}$ is the arc-length associated to the equatorial plane. Notice that $\chi_{ani} = 0$ corresponds to the isotropic constitutive equation for the active stress.

We found that enhanced circumferential cortex tension at the furrow exacerbated the local cell surface longitudinal stretching, decreasing the local apparent cell surface area compression (Fig. S15). An elevated circumferential cortical tension also reduced the accumulation of plasma membrane components at the division plane (Fig. S15). Our model results suggest that an elevated circumferential cortical stress could potentially have a negative impact on the completion of cytokinesis by increasing membrane tension at the intercellular bridge. This raises questions about the mechanical advantages of equatorial cortical tension anisotropy for promotion of cytokinesis. Note that anisotropic cortical tensions strongly influence the strength and polarization of cortical flows (42), with isotropic equatorial actomyosin stresses favoring cortical component recruitment at the equator. These effects are not included in our model.

#### Effects of plasma membrane mechanics on cell ingression.

Our primary focus has been on examining the impact of the actomyosin cortex on plasma membrane dynamics. Yet, it is important to acknowledge that the mechanical properties of the plasma membrane can reciprocally influence actomyosin and cleavage furrow ingression. Our model predicts that the strength of cortical flows and the movement of cortical components to the furrow is modulated by plasma membrane fluidity and membrane-cortex adhesion. Weaker flows and cortical recruitment are observed for gel-like membranes and high membrane-cortex adhesion, and stronger flows are observed for fluid-like membranes and low membrane-cortex adhesion (Fig. S16 and S7). This implies that cells with high membrane fluidity and/or low membrane-cortex adhesion experience elevated ring tension. Additionally, the membrane bending stiffness may contribute to cell constriction during furrow ingression. The emergence of membrane folding as the furrow ingresses can generate additional bending forces, potentially hindering cell constriction. Such bending forces are not a part of our force-balance model of cytokinesis.

### SUPPLEMENTARY NOTE 5: Limitations of the study

Our study has limitations that are important to highlight. Most notably, the exact implications of our findings to late cytokinesis and the final abscission of the daughter cells will require more investigation. The plasma membrane accumulation at the cleavage furrow will decrease over time due to membrane tension gradients along the pole-equator axis. However, the directed flow of plasma membrane components towards the poles can be weak, as it can be resisted by strong membrane-cortex drag forces (23). In agreement with this, the plasma membrane accumulation at the cleavage furrow did not fully disperse in our timelapse imaging, which extended ~20 min beyond furrow ingression (Fig. 3D). The full duration of cytokinesis in the L1210 cells, from the end of furrow ingression to final abscission, is 40±10 min (mean±SD) (9). Thus, membrane tension can remain low at the intercellular bridge during late cytokinesis due to the membrane accumulation detailed in this work.

Our live cell imaging experiments cannot quantify the total amounts of endo- and exocytosis. This prevents us from quantifying if exocytosis contributes significant amounts of new membrane to the cleavage furrow, as suggested before (43). Instead, our results verify that ample plasma membrane accumulation at the cleavage furrow takes place even in the absence of localized exocytosis. Our results are also predominantly obtained from a single model system, the L1210 cell line, and the magnitude of plasma membrane accumulation at the cleavage furrow due to exocytosis may vary significantly based on model system. Importantly, also the degree of plasma membrane accumulation at the cleavage furrow under *in vivo* conditions remains unclear as cell-extrinsic factors, such as cell-cell and cell-substrate interactions, can impact plasma membrane and actomyosin mechanics.

Our cytokinetic biophysical model also has some limitations. First, it does not consider cellular interactions with the extracellular environment. Cellular adhesion to the growth substrate and neighboring cells, for instance, can impact cytoskeletal organization during cytokinesis (44), and a strong adhesion to the environment could potentially cause additional resistance to cortical and plasma membrane flows towards the cleavage furrow. Nonetheless, even in cases where the plasma membrane resists these flows, the underlying mechanisms responsible for accumulating membrane at the cytokinetic furrow remain active. Second, our model does not account for active chiral forces which could significantly influence cortical flows and the mechanical deformation of the plasma membrane (45, 46). Third, our model assumes that both cortex-membrane and membrane lipid-protein drag coefficients remain spatially uniform and time-independent. However, we would anticipate the emergence of larger drag coefficients in the furrow region as cytokinesis progresses due to the accumulation of membrane proteins at the division plane, and the lower mobility of membrane components along folded membranes (47). These spatial variations may influence the dynamics of cortical flows, potentially suppressing cortical flows in the furrow region, especially during the later stages of ingression. Large variations in the estimated mobility values of plasma membrane components have been reported across cell types and physiological processes (13). Addressing these differences holds significant implications, not only for studying cell division but also for shedding light on other biological phenomena. Finally, our model does not account for the amplified assembly of actin filaments in the equatorial region (48). We assume that cortical association kinetics are spatially uniform, and recruitment of cortical components to the equatorial region during ring assembly is driven by cortical flows. Incorporating all these model components and performing cell-specific parameterization of model parameters is a potential opportunity for future experimental and theoretical studies on cytokinesis.

### SUPPLEMENTARY FIGURES

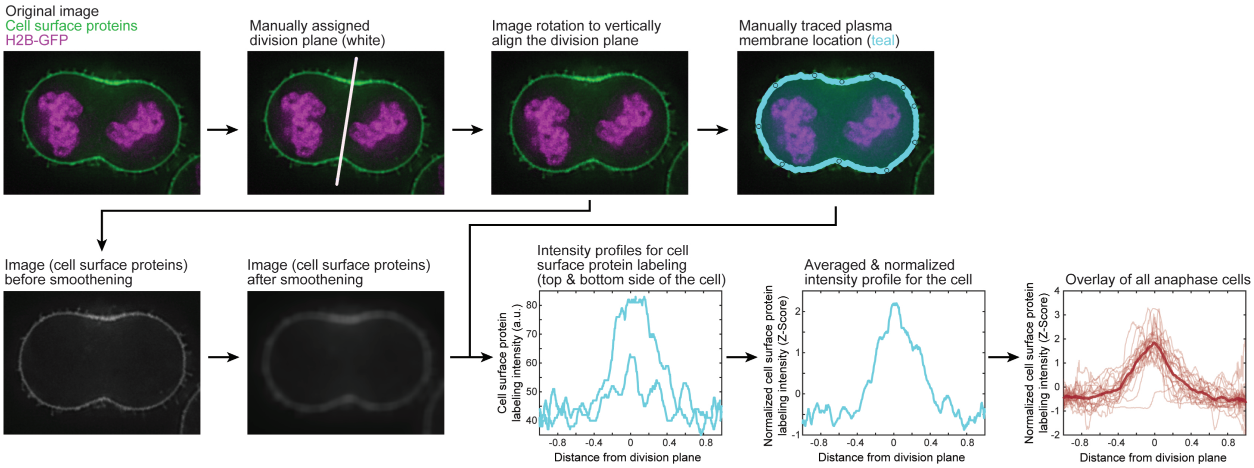

**Figure S1. Image analysis approach for local quantification of plasma membrane-specific labeling.** An example image of a cytokinetic L1210 cells expressing H2B-GFP (magenta) and labeled for surface protein content (green) and each step of the image analysis code. In the last step, light red lines depict individual cells and thick red line depicts the average of all cells. Data is the same as in Fig. 1B.

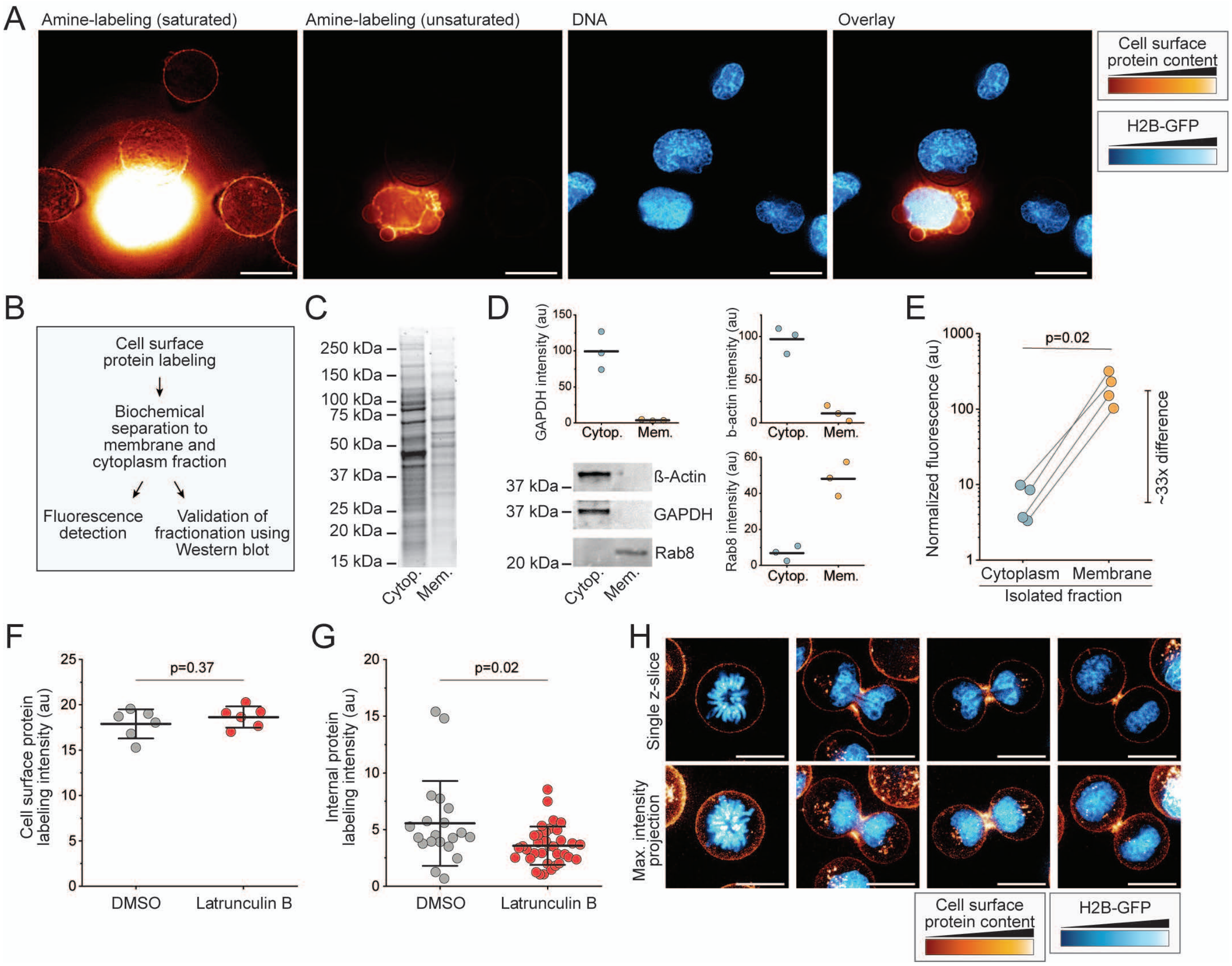

**Figure S2. Membrane protein labeling is independent of F-actin.**

**(A)** Representative images of L1210 cells expressing H2B-GFP (blue) and labeled for surface protein content using amine-reactive labeling chemistry (orange/yellow). One cell is blebbing and has lost membrane integrity, resulting in labeling of the cell's internal compartments and ~100-fold increased labeling intensity. N=8 FOVs that include dying cells. Scale bars denote 10 μm. **(B)** Experimental workflow. **(C)** Labeling of all proteins in the isolated cytoplasmic and membrane fractions after a separation using SDS PAGE. **(D)** Western blots of β-Actin, GAPDH, and Rab8 in the cytoplasmic and membrane fractions, along with quantifications from the blots (N=3 independent experiments). **(E)** Cell surface protein label fluorescence intensity in the isolated cytoplasmic and membrane fractions (N=4 independent experiments). **(F)** L1210 cell population average cell surface protein labeling intensity following a 2-hour treatment with DMSO or 2 μM Latrunculin B. Data depicts mean±SD of independent cultures (dots). N=2 independent experiments with 6 independent cultures. p-value was obtained using Welch's t-test. **(G)** L1210 cell internal labeling intensity following cell surface protein labeling and a treatment with DMSO or Latrunculin B. Data depicts mean±SD, dots depict single cells (n=20 and 35 cells for DMSO and Latrunculin B, respectively). p-value was obtained using Welch's t-test. **(H)** Representative images of mitotic and cytokinetic L1210 cells expressing H2B-GFP (blue) and labeled for surface protein content using thiol-reactive labeling chemistry (orange/yellow). N=2 independent experiments, n=17 cells. Thiol-reactive labeling displayed more intracellular accumulation of the label (endocytosis) than amine-reactive labeling (Fig. 1). Scale bars denote 10 μm.

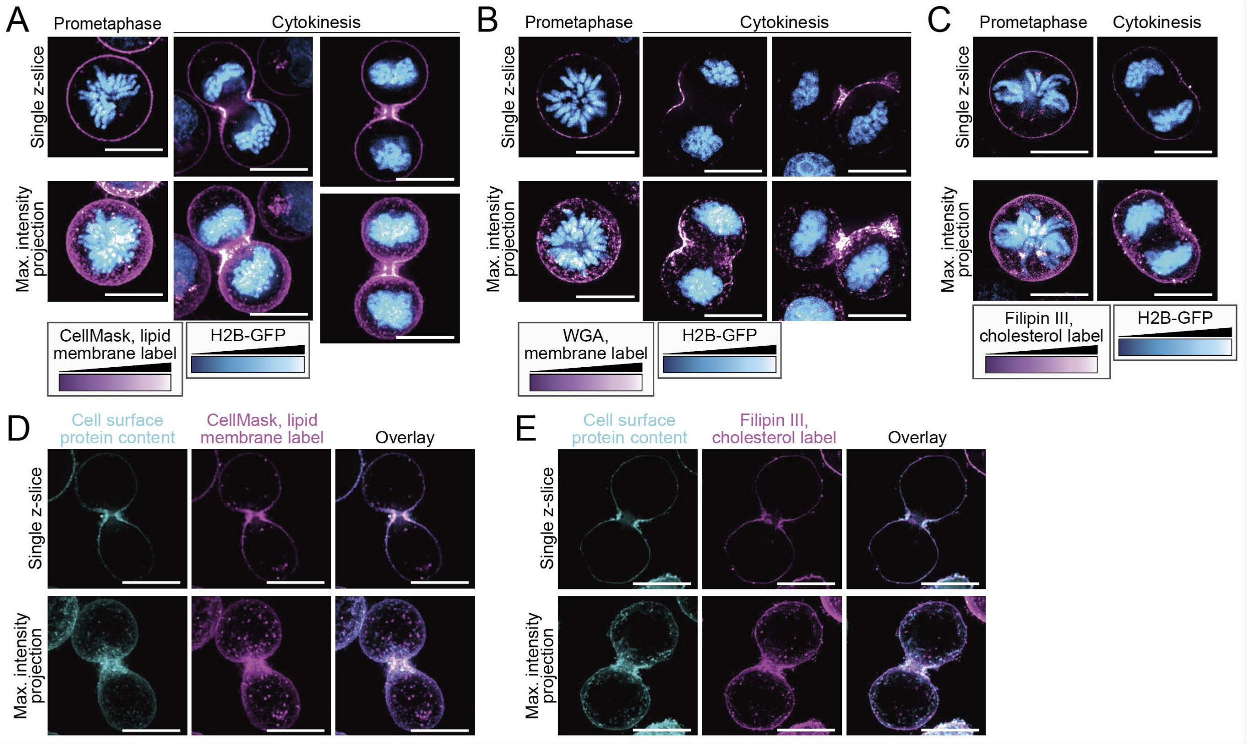

**Figure S3. Additional images of L1210 cell plasma membrane labels in cytokinesis.**
**(A-C)** Images of cytokinetic L1210 cells expressing H2B-GFP (blue, DNA) and labeled for plasma membrane (purple/white) using CellMask lipid label (panel A), WGA (panel B), or Filipin III (panel C). **(D)** Images of cytokinetic L1210 cells labeled using CellMask lipid label (purple) and cell surface protein label (teal). **(E)** Images of cytokinetic L1210 cells labeled using Filipin III cholesterol label (purple) and cell surface protein label (teal). Scale bars denote 10 μm.

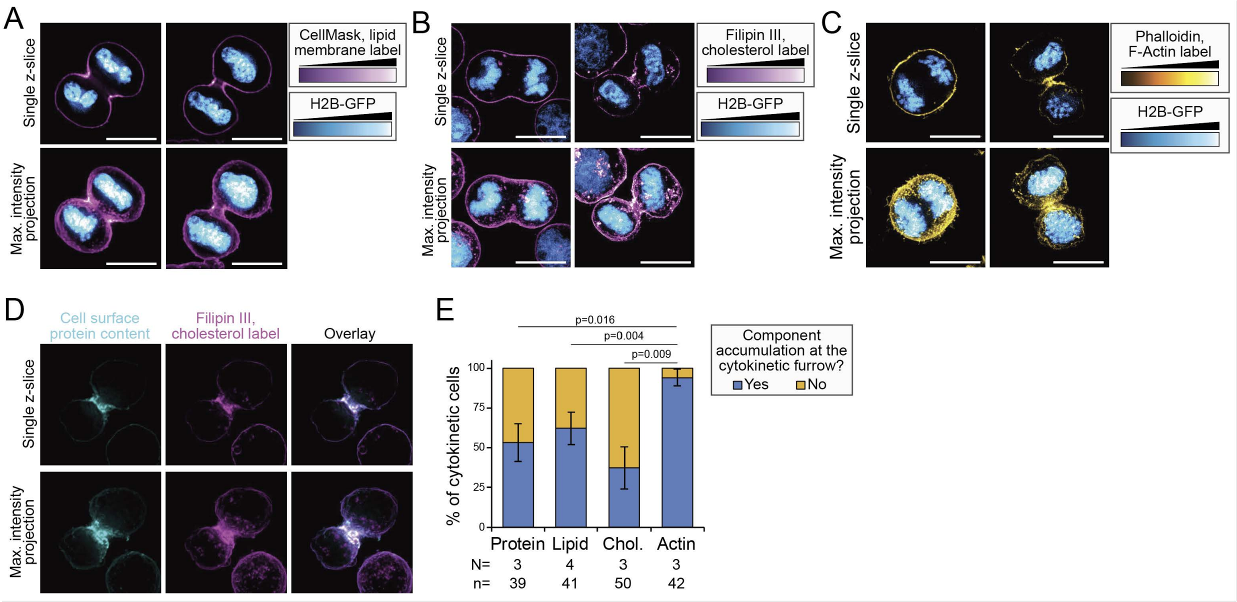

**Figure S4. Cytokinetic BaF3 cells display differential plasma membrane and F-actin dynamics.** **(A)** Representative images of live cytokinetic BaF3 cells expressing H2B-GFP (blue, DNA) and labeled for plasma membrane (purple/white) using CellMask lipid label. **(B)** Representative images of fixed cytokinetic BaF3 cells expressing H2B-GFP (blue, DNA) and labeled for cholesterol (purple/white) using Filipin III. **(C)** Representative images of fixed cytokinetic BaF3 cells expressing H2B-GFP (blue, DNA) and labeled for F-actin (yellow/white) using phalloidin. All scale bars denote 10 µm. **(D)** Representative images of a fixed cytokinetic BaF3 cell labeled for cell surface proteins (teal) and cholesterol (magenta). **(E)** Percentage of cytokinetic cells that exhibit the accumulation of plasma membrane proteins and lipids, and F-actin, at the cleavage furrow in BaF3 cells. Data depicts mean±SD of independent experiments. N depicts the number of independent experiments and n depicts the total number of cytokinetic cells. p-values were obtained using Welch's t-test. Protein labeling data is the same as in Fig. 1C.

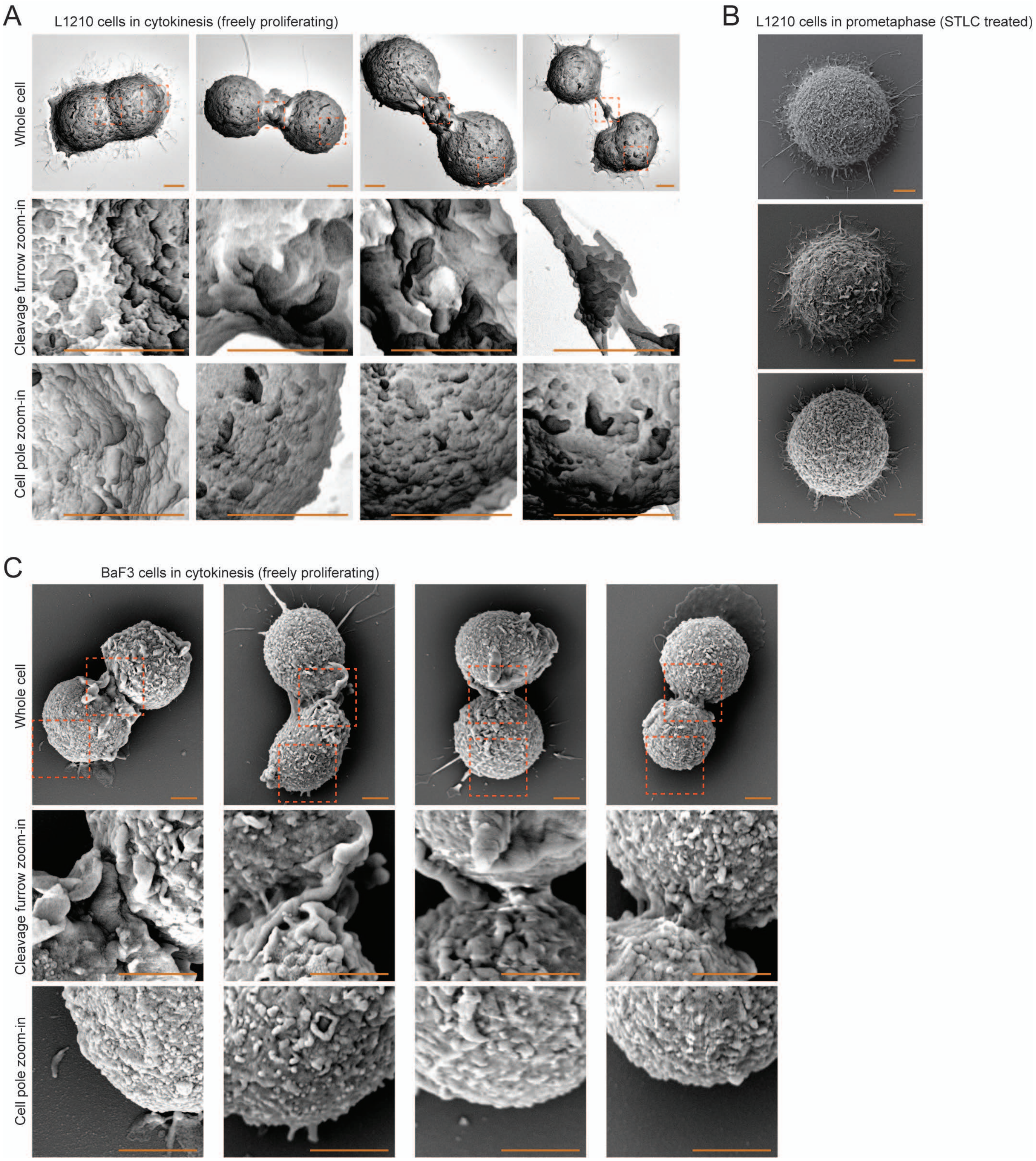

**Figure S5. SEM of plasma membrane folding in mitosis and cytokinesis.**
**(A)** Additional SEM images of fixed L1210 cells in cytokinesis, as shown in Fig. 2G. **(B)** Representative SEM images of L1210 cells arrested in prometaphase with 5 µM STLC for 10 hours (N=2 independent experiments, n=20 cells). **(C)** Representative SEM images of BaF3 cells in cytokinesis (N=2 independent experiment, n=16 cytokinetic cells). The two examples on left display cells with abundant membrane accumulation in cleavage furrow, while the two examples on right display less pronounced membrane accumulation. Scale bars denote 2 µm.

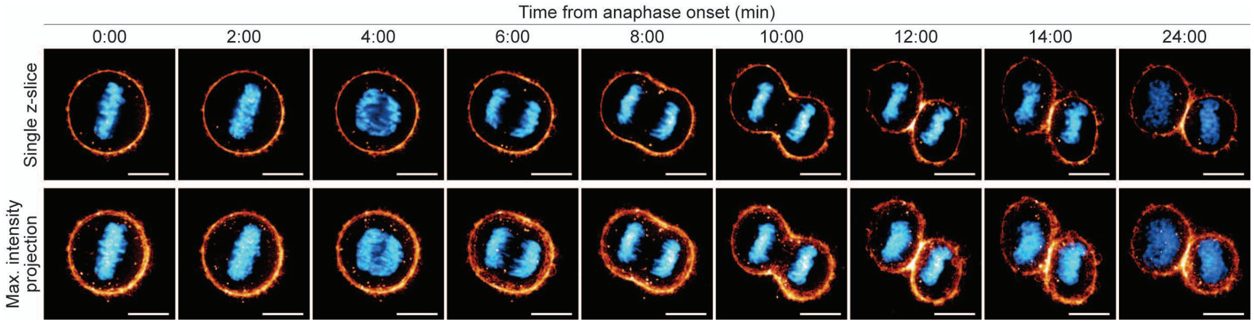

**Figure S6. Plasma membrane accumulation at the cleavage furrow is caused by movement of plasma membrane reservoirs on cell surface in Hela cells.**
Representative timelapse imaging of HeLa cells expressing H2B-GFP and labeled for cell surface protein content (orange/yellow). 13 out of 15 cytokinetic cells displayed the membrane accumulation at the furrow. N=4 independent experiments, n=15 cytokinetic cells. Scale bars denote 10 µm.

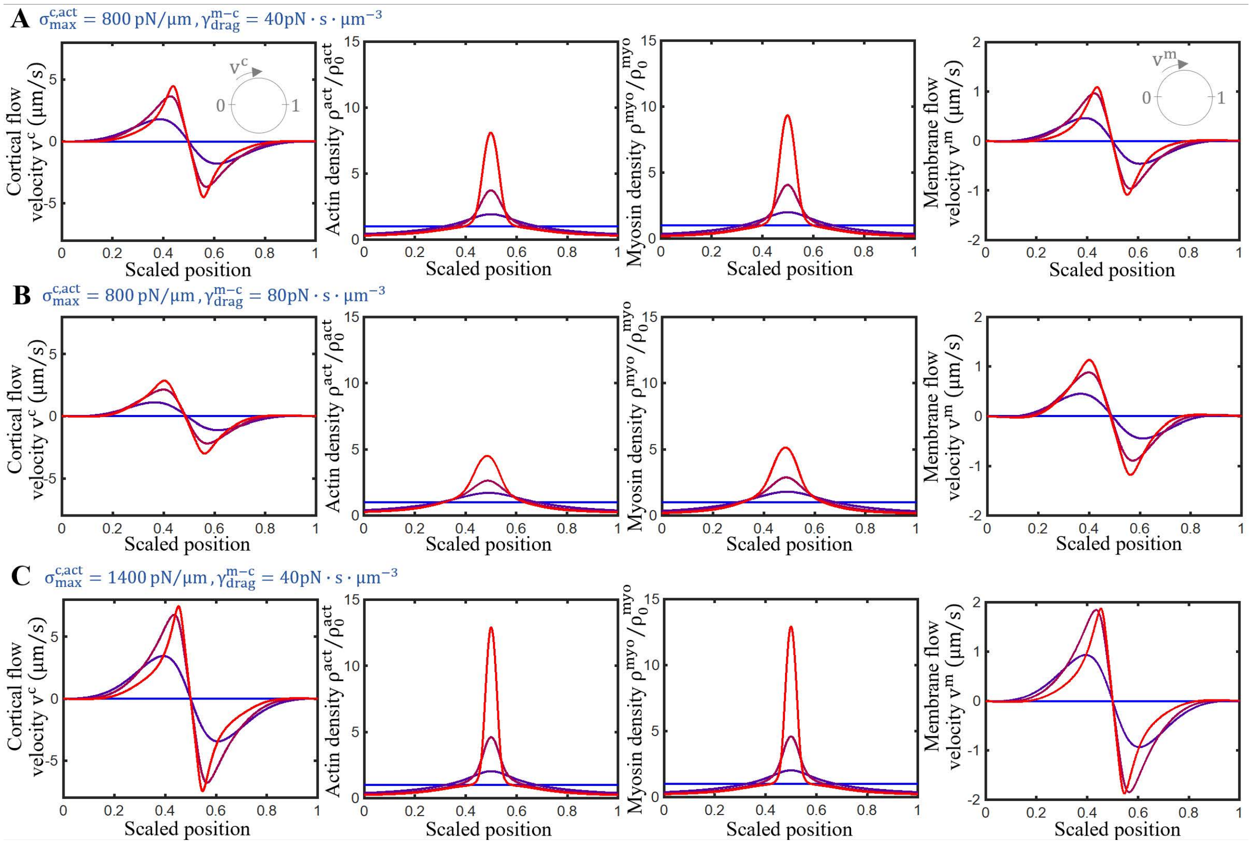

**Figure S7. Cortical and plasma membrane flows are directed towards the cell's equator.**
**(A-C)** Spatial distribution of in-plane cortical velocity, actin projected surface density, myosin projected surface density and in-plane membrane velocity for three different conditions. The strongest cortical and membrane flows are observed near the cell's equator (scaled position 0.5).

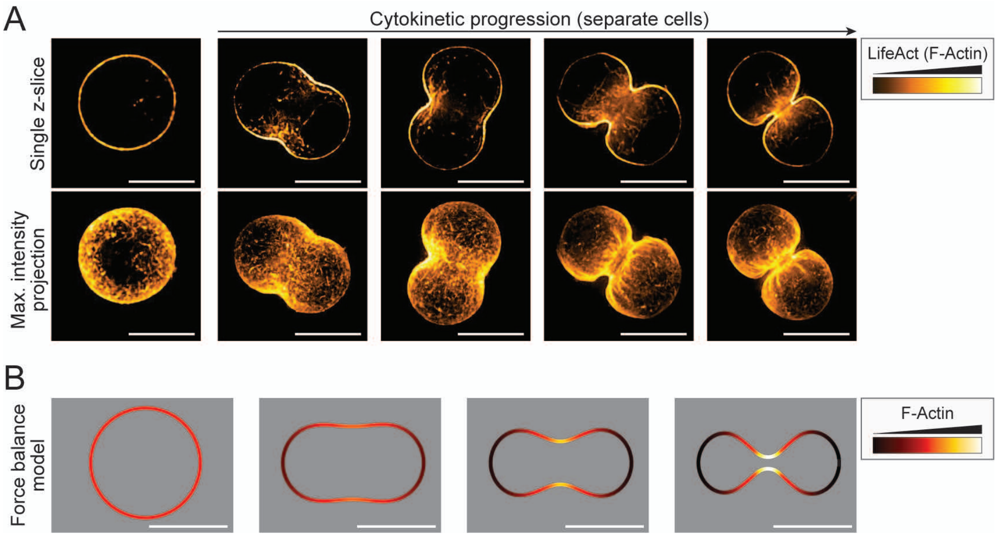

**Figure S8. Cell shapes and F-actin dynamics in live cells and in the cytokinesis model.**
**(A)** Representative images of live cytokinetic L1210 cells expressing the LifeAct F-actin sensor (n=14 cytokinetic cells). **(B)** Predictions of the cell shapes and local F-actin levels by the three-dimensional force balance model of cytokinesis. For simplicity images are presented in a single 2D plane. Scale bars denote 10 μm.

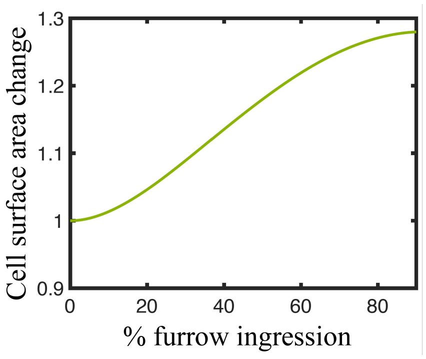

**Figure S9. Apparent cell surface area changes during cytokinesis.**
Changes in the total apparent cell surface area as a function of furrow ingression, as determined by the biophysical model of cytokinesis.

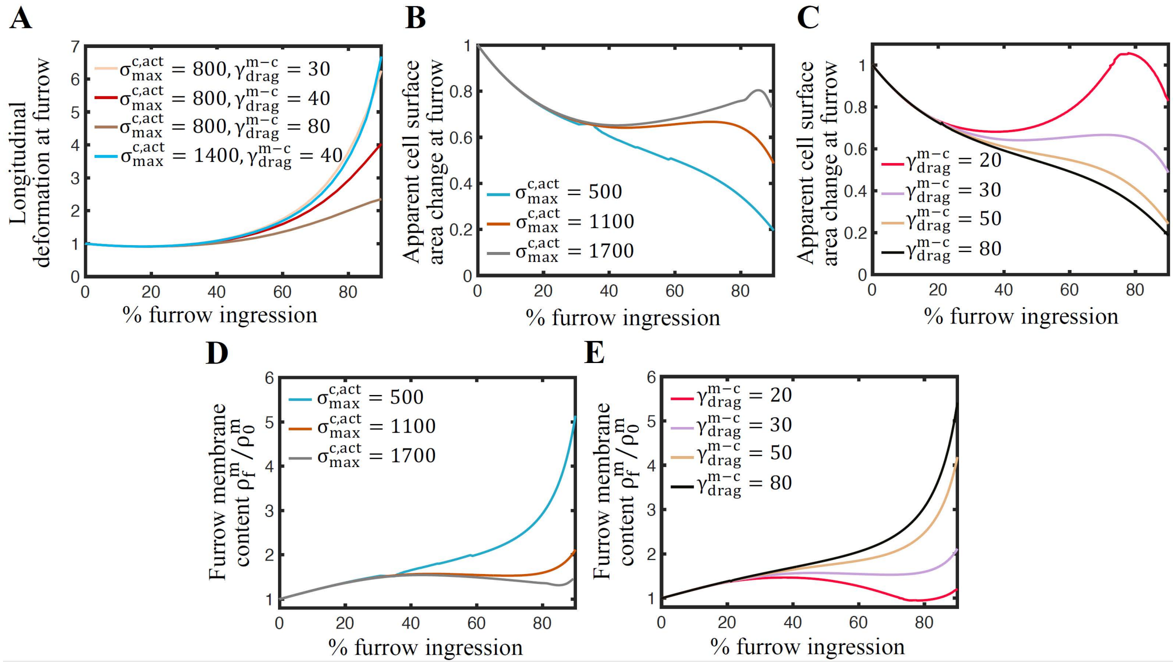

**Figure S10. Cells with low membrane fluidity, low membrane-cortex adhesion and high cortical tension do not exhibit plasma membrane accumulation at the cytokinetic furrow.**

**(A)** Longitudinal deformation at the furrow during furrow ingression for indicated levels of membrane-cortex drag coefficient ($\gamma_{\mathrm{drag}}^{\mathrm{m-c}}$, displayed in units of $\mathrm{pN \cdot s \cdot \mu m^{-3}}$) and cortical contractility ($\sigma_{\mathrm{max}}^{\mathrm{c,act}}$, displayed in units of $\mathrm{pN \cdot \mu m^{-1}}$). Cells with high cortical tension or low membrane-cortex adhesion exhibit enhanced longitudinal stretching. **(B-C)** Apparent cell surface area changes at the furrow (normalized to x=0) and **(D-E)** predicted plasma membrane accumulation at the furrow as furrow ingression progresses for cells with low plasma membrane fluidity ($\gamma_{\mathrm{drag}}^{\mathrm{m}} = 4000 \mathrm{pN \cdot s \cdot \mu m^{-3}}$) for different values of cortical contractility ($\sigma_{\mathrm{max}}^{\mathrm{c,act}}$) and membrane-cortex drag coefficient $\gamma_{\mathrm{drag}}^{\mathrm{m-c}}$. Parameter values: (A) $\gamma_{\mathrm{drag}}^{\mathrm{m}} = 100 \mathrm{pN \cdot s \cdot \mu m^{-3}}$, (B, D) $\gamma_{\mathrm{drag}}^{\mathrm{m-c}} = 30 \mathrm{pN \cdot s \cdot \mu m^{-3}}$, (C, E) $\sigma_{\mathrm{max}}^{\mathrm{c,act}} = 1100\ \mathrm{pN \cdot \mu m^{-1}}$.

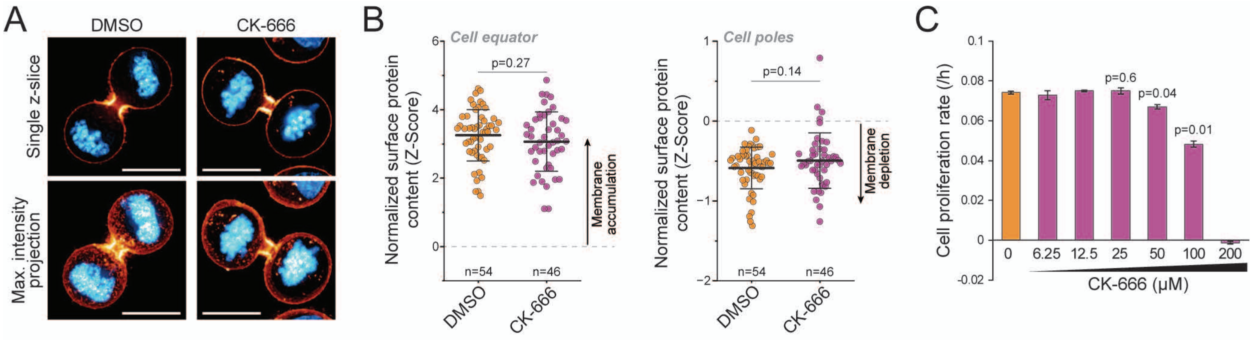

**Figure S11. CK-666 does not alter membrane accumulation at the cytokinetic furrow.**
**(A)** Representative images of cytokinetic L1210 cells expressing H2B-GFP (blue) and labeled for surface protein content (orange/yellow) after a 1 h treatment with DMSO or 50 μM CK-666. Scale bars denote 10 μm. **(B)** Quantifications of the surface protein content at the equatorial and polar regions in the samples shown in panel (A). Lines and whiskers depict mean±SD, dots depict individual cells, n depicts the number of cells (N=3 independent experiments). p-values were obtained using Welch's t-test. **(C)** Cell proliferation rate (doublings per hour) for indicated CK-666 concentrations. Bars and whiskers depict mean±SD. N=2 independent experiments. p-values reflect comparisons to DMSO control and were obtained using Welch's t-test.

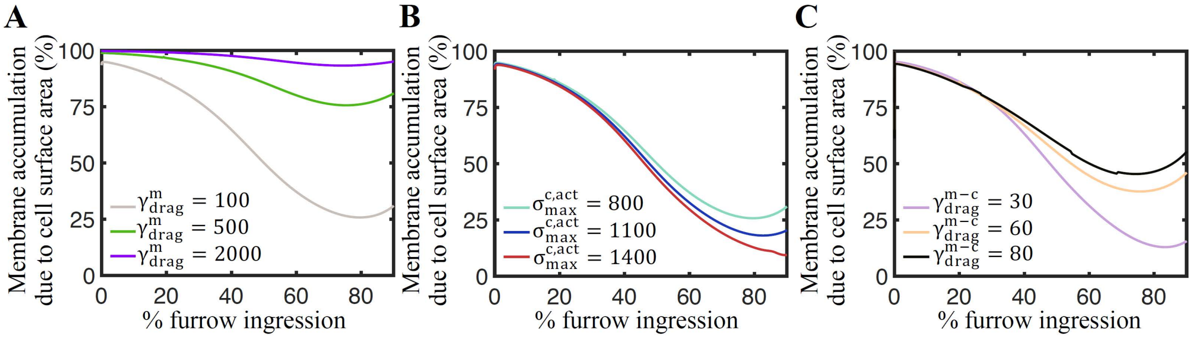

**Figure S12. Plasma membrane fluidity and membrane-cortex adhesion impact plasma membrane accumulation at the cytokinetic furrow.**

**(A-C)** Percentage of plasma membrane accumulation at the cleavage furrow attributed to local changes in cell surface area in contrast to cortex-drag forces. Data is displayed for different values of the membrane lipid-protein drag coefficient (A), cortical contractility (B) and membrane-cortex drag coefficient (C). High membrane fluidity, high cortical tension and low membrane-cortex adhesion increase the relative contribution to membrane accumulation due to cortex-drag forces.

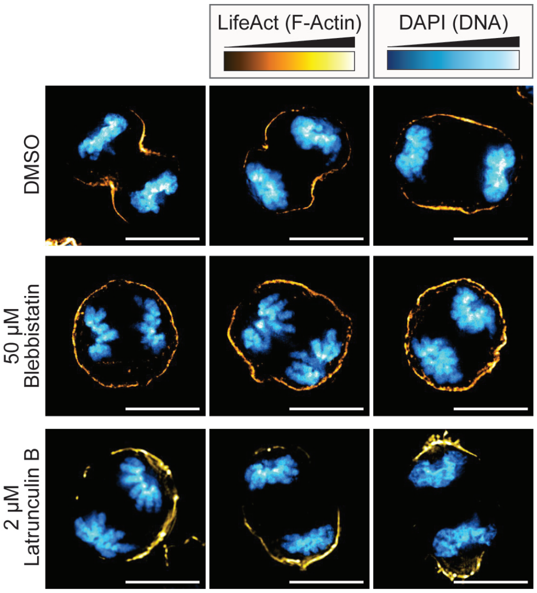

**Figure S13. F-actin distribution following drug treatments in anaphase L1210 cells.**
Representative images of fixed L1210 cells expressing the LifeAct F-actin sensor (yellow) and labeled for DNA (blue). The cells were treated with DMSO, 50 µM (-)-Blebbistatin or 2 µM Latrunculin B for 2 hours. N=3 independent experiments, n=23, 18 and 30 cells for DMSO, Blebbistatin and Latrunculin B treatments, respectively. Scale bars denote 10 µm.

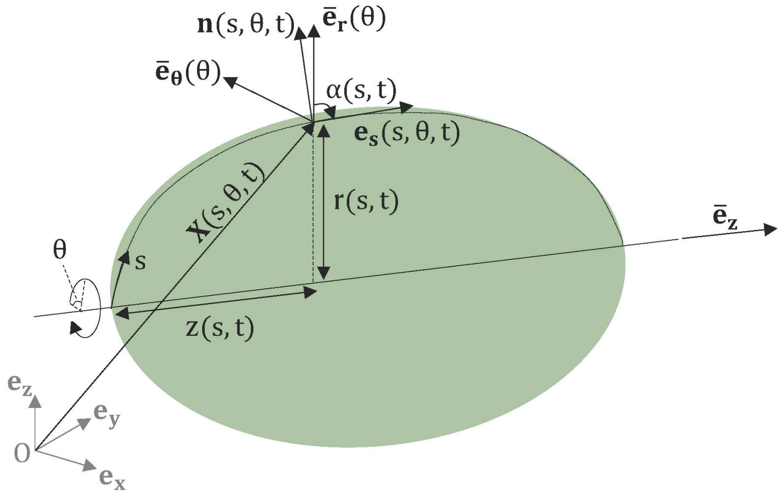

**Figure S14. Parameterization of the cell surface.**

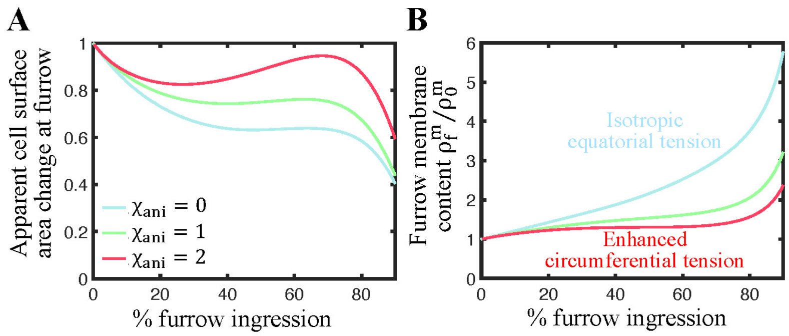

**Figure S15. Enhanced circumferential tension decreases plasma membrane accumulation at the cytokinetic furrow.**
**(A)** Changes in the apparent cell surface area ($A_f$) as a function of furrow ingression for three values of the anisotropic tension parameter. Enhanced circumferential cortical tension at the furrow decreases the degree of local cell surface area compression. **(B)** Furrow membrane content as ingression progresses for the three different anisotropic tension parameter values. Enhanced circumferential tension decreases the accumulation of plasma membrane components at the division plane. Parameter value: $h_{ani} = 2\mu m$.

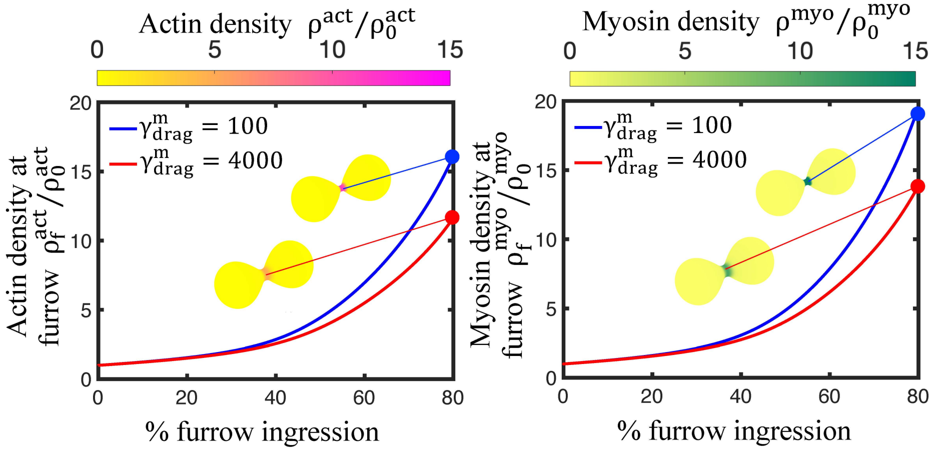

**Figure S16. Recruitment of cortical actin and myosin is predicted to be enhanced in cells with high plasma membrane fluidity.**

Predicted actin and myosin projected densities at the cytokinetic furrow as ingression progresses for two different values of the membrane lipid-protein drag coefficient. Insets show snapshot of the predicted actin and myosin cortical distributions at 80% of furrow ingression. Parameter values: $\gamma_{drag}^{m-c} = 30\ \mathrm{pN \cdot s \cdot \mu m^{-3}}$, $\sigma_{max}^{c,act} = 800\ \mathrm{pN \cdot \mu m^{-1}}$.

### LEGENDS FOR SUPPLEMENTARY MOVIES

**Movie S1. Plasma membrane accumulates at the cleavage furrow via movement on the cell surface.** A representative movie of dividing L1210 cells expressing H2B-GFP (blue) and labeled for cell surface protein content (orange/yellow). Each timepoint is a maximum intensity projection of 11 z-layers. N=3 independent experiments, n=14 cytokinetic cells. Scale bar depicts 10 μm.

**Movie S2. Apparent cell surface area changes and longitudinal stretching of the cell surface in the force balance model of cytokinesis.**
Cytokinesis was modelled using a membrane-cortex drag coefficient ($\gamma_{\mathrm{drag}}^{\mathrm{m-c}}$) of 40 $\mathrm{pN \cdot s \cdot \mu m^{-3}}$, cortical contractility ($\sigma_{\mathrm{max}}^{\mathrm{c,act}}$) of 800 $\mathrm{pN \cdot \mu m^{-1}}$, and membrane lipid-protein drag coefficient ($\gamma_{\mathrm{drag}}^{\mathrm{m}}$) of 100 $\mathrm{pN \cdot s \cdot \mu m^{-3}}$. These are the standard modeling values used throughout our manuscript.

**Movie S3. Apparent cell surface area changes and longitudinal stretching of the cell surface in the force balance model of cytokinesis, when using alternative model parameters.**
Cytokinesis was modelled using a membrane-cortex drag coefficient ($\gamma_{\mathrm{drag}}^{\mathrm{m-c}}$) of 60 $\mathrm{pN \cdot s \cdot \mu m^{-3}}$, cortical contractility ($\sigma_{\mathrm{max}}^{\mathrm{c,act}}$) of 1200 $\mathrm{pN \cdot \mu m^{-1}}$, and membrane lipid-protein drag coefficient ($\gamma_{\mathrm{drag}}^{\mathrm{m}}$) of 2000 $\mathrm{pN \cdot s \cdot \mu m^{-3}}$.

**Movie S4. Plasma membrane dynamics predicted by the force balance model of cytokinesis.**
Cytokinesis was modelled using a membrane-cortex drag coefficient ($\gamma_{\mathrm{drag}}^{\mathrm{m-c}}$) of 40 $\mathrm{pN \cdot s \cdot \mu m^{-3}}$, cortical contractility ($\sigma_{\mathrm{max}}^{\mathrm{c,act}}$) of 800 $\mathrm{pN \cdot \mu m^{-1}}$, and membrane lipid-protein drag coefficient ($\gamma_{\mathrm{drag}}^{\mathrm{m}}$) of 100 $\mathrm{pN \cdot s \cdot \mu m^{-3}}$. These are the standard modeling values used throughout our manuscript.

**Movie S5. Plasma membrane dynamics predicted by the force balance model of cytokinesis, when using alternative model parameters.**
Cytokinesis was modelled using a membrane-cortex drag coefficient ($\gamma_{\mathrm{drag}}^{\mathrm{m-c}}$) of 60 $\mathrm{pN \cdot s \cdot \mu m^{-3}}$, cortical contractility ($\sigma_{\mathrm{max}}^{\mathrm{c,act}}$) of 1200 $\mathrm{pN \cdot \mu m^{-1}}$, and membrane lipid-protein drag coefficient ($\gamma_{\mathrm{drag}}^{\mathrm{m}}$) of 2000 $\mathrm{pN \cdot s \cdot \mu m^{-3}}$.
